# Graph theory approaches to functional network organization in brain disorders: A critique for a brave new small-world

**DOI:** 10.1101/243741

**Authors:** Michael N. Hallquist, Frank G. Hillary

**Affiliations:** Department of Psychology, Pennsylvania State University, University Park, PA; Social Life and Engineering Sciences Imaging Center, University Park, PA; Department of Neurology, Hershey Medical Center, Hershey, PA

**Author notes:** Corresponding author: Frank G. Hillary, Department of Psychology, Penn State University, 140 Moore Building, University Park, PA 16802.

**Keywords:** graph theory, brain disorders, network neuroscience, proportional thresholding

## Abstract

Over the past two decades, resting-state functional connectivity (RSFC) methods have provided new insights into the network organization of the human brain. Studies of brain disorders such as Alzheimer’s disease or depression have adapted tools from graph theory to characterize differences between healthy and patient populations. Here, we conducted a review of clinical network neuroscience, summarizing methodological details from 106 RSFC studies. Although this approach is prevalent and promising, our review identified four challenges. First, the composition of networks varied remarkably in terms of region parcellation and edge definition, which are fundamental to graph analyses. Second, many studies equated the number of connections across graphs, but this is conceptually problematic in clinical populations and may induce spurious group differences. Third, few graph metrics were reported in common, precluding meta-analyses. Fourth, some studies tested hypotheses at one level of the graph without a clear neurobiological rationale or considering how findings at one level (e.g., global topology) are contextualized by another (e.g., modular structure). Based on these themes, we conducted network simulations to demonstrate the impact of specific methodological decisions on case-control comparisons. Finally, we offer suggestions for promoting convergence across clinical studies in order to facilitate progress in this important field.

## Introduction

Efforts to characterize a “human connectome” have brought sweeping changes to functional neuroimaging research, with many investigators transitioning from indices of local brain activity to measures of inter-regional communication (Friston, 2011). The broad goal of this conceptual revolution is to understand the brain as a functional network whose coordination is responsible for complex behaviors (Biswal et al., 2010). The prevailing approach to studying functional connectomes involves quantifying coupling of the intrinsic brain activity among regions. In particular, resting-state functional connectivity (RSFC) methods (Biswal, Yetkin, Haughton, & Hyde, 1995) focus on inter-regional correspondence in low-frequency oscillations of the BOLD signal (approximately 0.01-0.12 Hz).

Work over the past two decades has demonstrated the value of RSFC approaches for mapping functional network organization, including the identification of separable brain subnetworks (Biswal et al., 2010; Laird et al., 2009; Power et al., 2011; Smith et al., 2009; van den Heuvel & Hulshoff Pol, 2010). Because RSFC methods do not require the study-specific designs and cognitive burden associated with task-based fMRI studies, RSFC data are simple to acquire and have been used in hundreds of studies of human brain function. Nevertheless, there are numerous methodological challenges, including concerns about the quality of RSFC data (Power et al., 2014) and the effect of data processing on substantive conclusions (Ciric et al., 2017; Hallquist, Hwang, & Luna, 2013; Shirer, Jiang, Price, Ng, & Greicius, 2015).

RSFC studies of brain injury or disease typically examine differences in the functional connectomes of a clinical group (e.g., Parkinson’s disease) compared to a matched control group. There are specific methodological and substantive considerations that apply to RSFC studies of brain disorders. For example, differences in the overall level of functional connectivity between patient and control groups could lead to differences in the number of spurious connections in network analyses, potentially obscuring meaningful group comparisons (van den Heuvel et al., 2017). Likewise, there is increasing concern in the clinical neurosciences that an unacceptably small percentage of findings are replicable (Müller et al., 2017). Such concerns echo the growing emphasis on open, reproducible practices in neuroimaging more generally (Poldrack et al., 2017).

In this paper, we review the current state of graph theory approaches to RSFC in the clinical neurosciences. Based on key themes in this literature, we conducted two network simulations to demonstrate the pitfalls of specific analysis decisions that have particular relevance to case-control studies. Finally, we provide recommendations for best practices to promote comparability across studies.

Our review does not directly address many important methodological issues that are active areas of investigation. For example, detecting and correcting motion-related artifacts remains a central challenge in RSFC studies (Ciric et al., 2017; Dosenbach et al., 2017) that is especially problematic in clinical and developmental samples (Greene, Black, & Schlaggar, 2016; Van Dijk, Sabuncu, & Buckner, 2012). The length of resting-state scans (in terms of time, number of measurements, and number of sessions) also influences the reliability of functional connectivity estimates (Birn et al., 2013; Gonzalez-Castillo, Chen, Nichols, & Bandettini, 2017) and likely has downstream consequences on network analyses (Abrol et al., 2017; Yang et al., 2012). Finally, brain parcellation — defining the number and form of brain regions — is one of the most important influences on the composition of RSFC networks. There are numerous parcellation approaches, including anatomical atlases, functional boundary mapping, and data-driven algorithms based on voxelwise BOLD time series (Goñi et al., 2014; Honnorat et al., 2015). In order to focus on RSFC graph theory research in the clinical literature, where relevant, we refer readers to more focused treatments of important issues that are beyond the scope of this paper.

### A Literature Review of Clinical Network Neuroscience Studies

Graph theory is a branch of discrete mathematics that has been applied in numerous studies of brain networks, both structural and functional. A *graph* is a collection of objects, called *vertices* or *nodes*; the pairwise relationships among nodes are called *edges* or *links* (Newman, 2010). Graphs composed of RSFC estimates among regions provide a window into the intrinsic connectivity patterns in the human brain. Figure 1 provides a simple schematic of the most common graph theory constructs and metrics reported in RSFC studies. For more comprehensive reviews of graph theory applications in network neuroscience, see Bullmore and Sporns (2009,2012), or Fornito, Zalesky, and Bullmore (2016).

**Figure 1.**
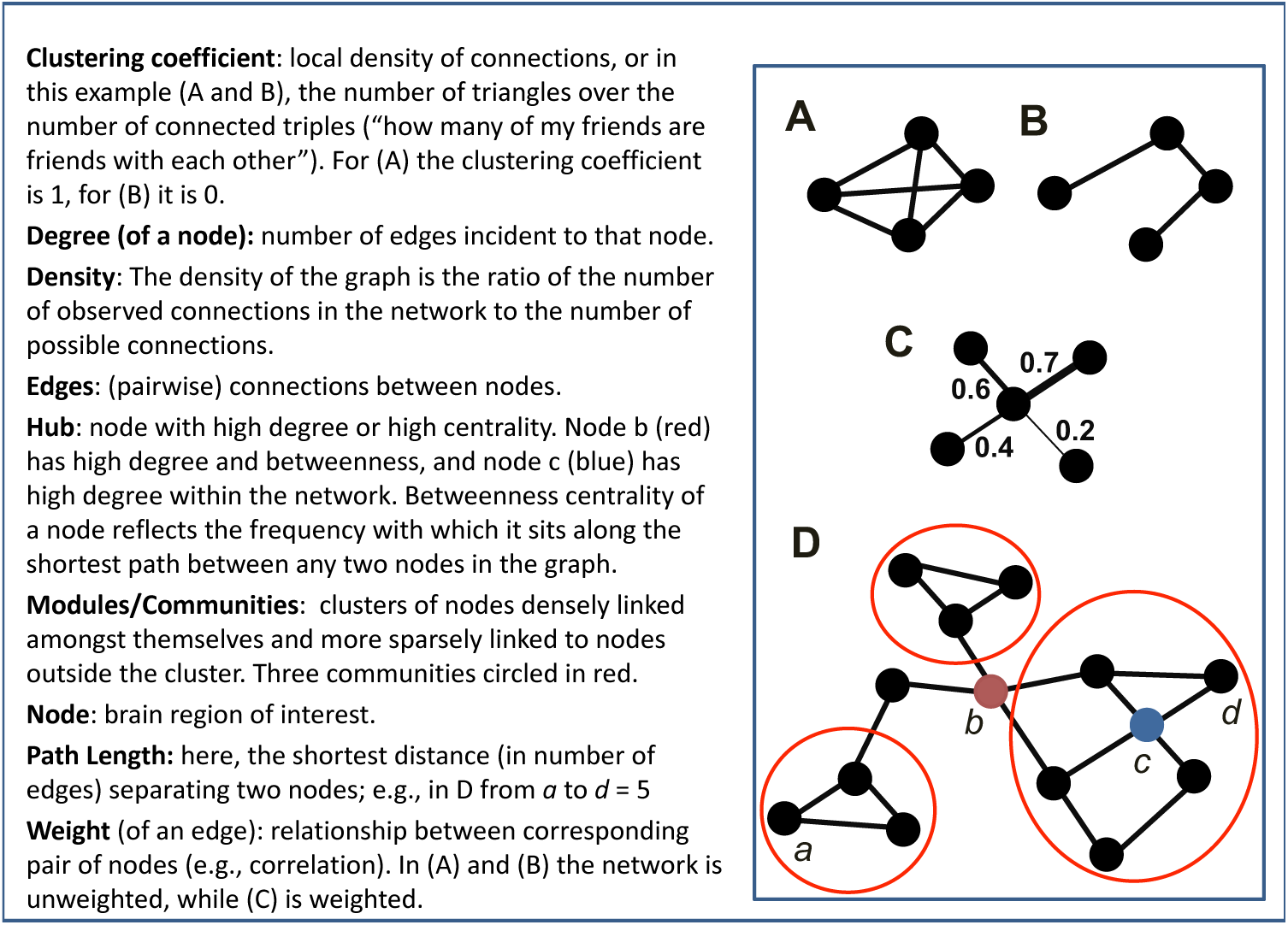
A toy graph and related graph theory terminology. Adapted with permission from Hillary & Grafman, 2017.

The goal of our literature search was to obtain a representative cross-section of graph-theoretic RSFC studies spanning neurological and mental disorders. We focused on functional connectivity (FC) as opposed to structural connectivity, where a distinct set of methodological critiques are likely relevant. Also, we reviewed fMRI studies only, excluding electroencephalography (EEG) and magnetoencephalography (MEG). Although there are important advantages of EEG/MEG in some respects (Papanicolaou et al., 2017), we focused on fMRI in part because the vast majority of clinical RSFC studies have used this modality. In addition, there are fMRI-specific considerations for network definition and spatial parcellation in RSFC studies. We conducted two related searches of the PubMed database (http://www.ncbi.nlm.nih.gov/pubmed) to identify articles focusing on graph-theoretic approaches to RSFC in mental and neurological disorders (for details on the queries, see Methods).

These searches were run in April 2016 and resulted in 626 potential papers for review (281 from neurological query, 345 from mental disorder query). Studies were excluded if they were reviews, case studies, animal studies, methodological papers, used electrophysiological methods (e.g., EEG or MEG), reported only structural imaging, or did not focus on brain disorders (e.g., healthy brain functioning, normal aging). After exclusions, these two searches yielded distinct 106 studies included in the review (see Table 1). A full listing of all studies reviewed is provided in the Supplementary Materials.

**Table 1.**
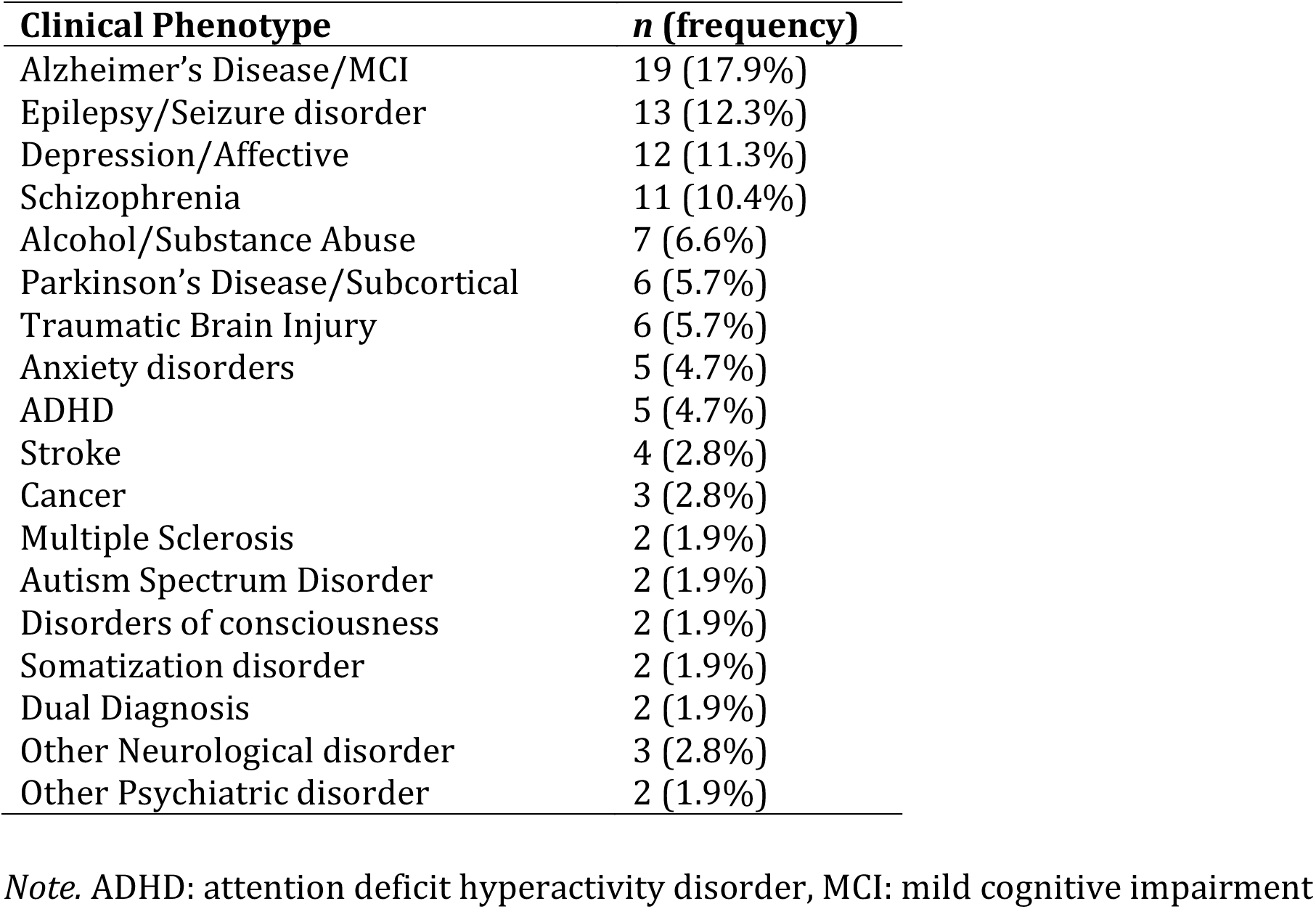
Clinical disorders represented in the review of 106 clinical network neuroscience studies

| **Clinical Phenotype** | ***n* (frequency)** |
| --- | --- |
| Alzheimer's Disease/MCI | 19 (17.9%) |
| Epilepsy/Seizure disorder | 13 (12.3%) |
| Depression/Affective | 12 (11.3%) |
| Schizophrenia | 11 (10.4%) |
| Alcohol/Substance Abuse | 7 (6.6%) |
| Parkinson's Disease/Subcortical | 6 (5.7%) |
| Traumatic Brain Injury | 6 (5.7%) |
| Anxiety disorders | 5 (4.7%) |
| ADHD | 5 (4.7%) |
| Stroke | 4 (2.8%) |
| Cancer | 3 (2.8%) |
| Multiple Sclerosis | 2 (1.9%) |
| Autism Spectrum Disorder | 2 (1.9%) |
| Disorders of consciousness | 2 (1.9%) |
| Somatization disorder | 2 (1.9%) |
| Dual Diagnosis | 2 (1.9%) |
| Other Neurological disorder | 3 (2.8%) |
| Other Psychiatric disorder | 2 (1.9%) |

*Note.* ADHD: attention deficit hyperactivity disorder, MCI: mild cognitive impairment

Below, we outline important general themes from the literature review (our major concerns are summarized in Box 1), including the heterogeneity of data analytic approaches across graph theoretical studies. We then turn our focus to two critical issues with important implications for interpreting network analyses in case-control studies: 1) network thresholding (i.e., determining how to define a “connection” from a continuous measure of FC) and 2) matching the hypothesis to the level of inquiry in the graph. For each of these issues, we offer network simulations to illustrate the importance of these issues for case-control comparisons.

##### Box 1: Major challenges in graph-theoretic studies of functional connectivity in brain disorders

1. There is substantial heterogeneity in defining the nodes and edges of graphs across resting-state studies that largely precludes quantitative comparisons and formal meta-analyses. Key sources of heterogeneity include: brain parcellations, functional connectivity metrics (e.g., full versus partial correlation), handling of negative FC estimates, and preprocessing decisions (e.g., global signal regression). Furthermore, few graph metrics were reported in common, further undermining an examination of similarities across studies and disorders.
2. Many studies use proportional thresholding to convert a continuous functional connectivity metric (e.g., Pearson correlation) to a binary edge in the graph while controlling for the total number of edges. This approach may obscure the effects of brain pathology in case-control designs, where clinical groups may have changes in the number and strength of functional connections. Proportional thresholding may also identify spurious group differences when groups differ in the connectivity strength of selected brain regions.
3. The link between many graph metrics and neurobiology has been more commonly presumed than established (e.g., neural efficiency).
4. The alignment between study hypotheses and specific graph analyses was often unclear. Moreover, many studies have not considered how findings at one level (e.g., global topology) are contextualized by another (e.g., modular structure).

### Creating Comparable Networks in Clinical Samples

#### Defining nodes in functional brain networks

Graph theory analyses of RSFC data fundamentally depend on the definition of nodes (i.e., brain regions) and edges (i.e., the quantification of functional connectivity). For network analyses to reveal new insights into clinical phenomena, investigators must choose a parcellation scheme that robustly samples the regions and networks of interest. Our literature review revealed that 76% of studies defined graphs based on comprehensive parcellations (i.e., sampling most or all of the brain), whereas 24% analyzed connectivity in targeted sub-networks (e.g., motor regions only; Table 2a). Furthermore, we found substantial heterogeneity in parcellations, ranging from 10 to 67632 nodes (Mode = 90; *M* = 1129.2; *SD* = 7035.9). In fact, whereas 25% of studies had 90 nodes (most of these used the AAL atlas; Tzourio-Mazoyer et al., 2002), the frequency of all other parcellations fell below 5%, resulting in at least 50 distinct parcellations in 106 studies.

**Table 2.**
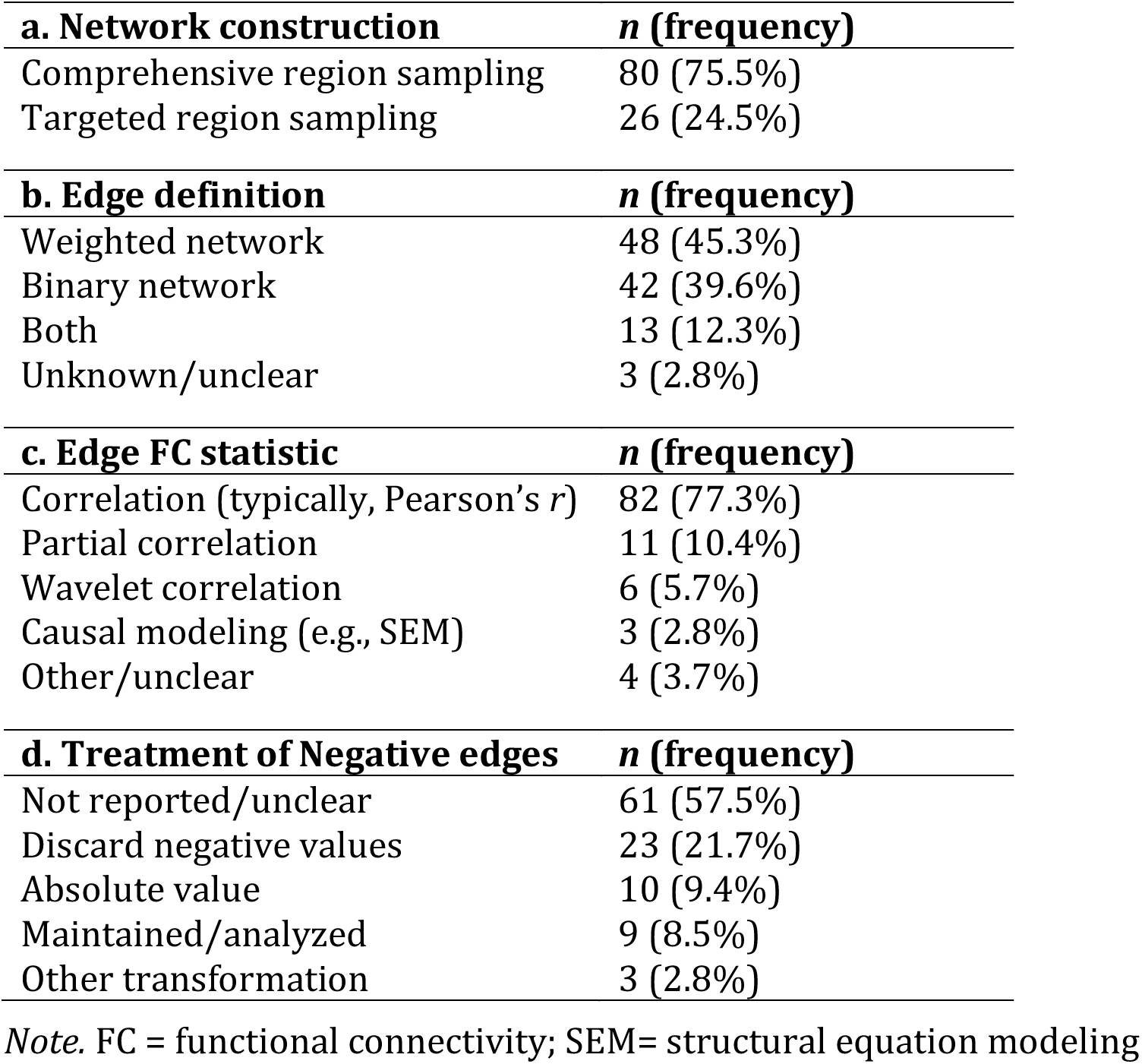
Network construction and edge definition

| **a. Network construction** | ***n* (frequency)** |
| --- | --- |
| Comprehensive region sampling | 80 (75.5%) |
| Targeted region sampling | 26 (24.5%) |
| **b. Edge definition** | ***n* (frequency)** |
| Weighted network | 48 (45.3%) |
| Binary network | 42 (39.6%) |
| Both | 13 (12.3%) |
| Unknown/unclear | 3 (2.8%) |
| **c. Edge FC statistic** | ***n* (frequency)** |
| Correlation (typically, Pearson's *r*) | 82 (77.3%) |
| Partial correlation | 11 (10.4%) |
| Wavelet correlation | 6 (5.7%) |
| Causal modeling (e.g., SEM) | 3 (2.8%) |
| Other/unclear | 4 (3.7%) |
| **d. Treatment of Negative edges** | ***n* (frequency)** |
| Not reported/unclear | 61 (57.5%) |
| Discard negative values | 23 (21.7%) |
| Absolute value | 10 (9.4%) |
| Maintained/analyzed | 9 (8.5%) |
| Other transformation | 3 (2.8%) |

*Note.* FC = functional connectivity; SEM= structural equation modeling

Although it may seem self-evident, it bears mentioning that several popular parcellation schemes provide a broad, but not complete, sampling of functional brain regions. For example, recent parcellations based on the cortical surface of the brain (e.g.,Glasser et al., 2016; Gordon et al., 2016) have provided a new level of detail on functional boundaries in the cortex. Yet if a researcher is interested in cortical-subcortical connectivity, it is crucial that the parcellation be extended to include all relevant regions.

There are advantages and challenges to every parcellation approach (e.g.,Honnorat et al., 2015; Power et al., 2011); here, we focus on two specific concerns. First, a goal of most clinical network neuroscience studies is to describe group differences in whole-brain connectivity patterns that are reasonably robust to the graph definition. Thus, investigators may wish to use at least two parcellations in the same data set to determine if the findings are parcellation-dependent. Because of the fundamental difficulty in comparing unequal networks (van Wijk, Stam, & Daffertshofer, 2010), one would not expect identical findings. In particular, global topological features such as efficiency or characteristic path length may vary by parcellation, but other features such as modularity and hub architecture are likely to be more robust. Applying multiple parcellations to the same dataset increases the number of analyses multiplicatively, as well as the need to reconcile inconsistent findings. Cassidy and colleagues (2018) recently demonstrated that persistent homology, a technique from topological analysis, has the potential to quantify the similarity of individual functional connectomes across different parcellations. We believe that such directions hold promise for supporting reproducible findings in clinical RSFC studies (Craddock, James, Holtzheimer, Hu, & Mayberg, 2012; Roy et al., 2017; Schaefer et al., in press).

Second, differences in parcellation fundamentally limit the ability to compare studies, both descriptively and quantitatively. As noted above, our review revealed substantial heterogeneity in parcellation schemes across studies of brain disorders. Extending our concern about parcellation dependence, such heterogeneity makes it impossible to ascertain whether differences between two studies of the same clinical population are an artifact of the graph definition or a meaningful finding. Moreover, whereas meta-analyses of structural MRI and task-based fMRI studies have become increasingly popular (e.g.,Goodkind et al., 2015; Müller et al., 2017), such analyses are not currently possible in graph-theoretic studies in part because of differences in parcellation. To resolve this issue, we encourage scientists to report results for a field-standard parcellation, a point we elaborate in our recommendations below. We also believe there is value in investigators having freedom to test and compare additional parcellations that may highlight specific findings.

#### Defining edges in functional brain networks

Parcellation defines the nodes comprising a graph, but an equally important decision is how to define functional connections among nodes (i.e., the edges). The vast majority (77%) of the studies reviewed used bivariate correlation, especially Pearson or Spearman, as the measure of FC. In Critical Issue 1 below, we consider how FC estimates are thresholded in order to defines edges as present or absent in binary graphs.

##### Quantifying functional connectivity

The statistical measure of FC has important implications for network density and the interpretation of relationships among brain regions. The prevailing application of marginal association (typically, bivariate correlation) does not separate the (statistically) direct connectivity between two regions from indirect effects attributable to additional regions. By contrast, most conditional association methods (e.g., partial correlation) rely on inverting the covariance matrix among all regions, thereby removing common variance and defining edges based on unique connectivity between two regions (Smith et al., 2011).

At the present time, there is no clear resolution on whether FC should be based on marginal or conditional association (for useful reviews, see Fallani, Richiardi, Chavez, & Achard, 2014; Varoquaux & Craddock, 2013). Nevertheless, we wish to highlight that as the average full correlation increases within a network, partial correlation values, on average, must decrease. Furthermore, partial and full correlations reveal fundamentally different graph topologies in resting-state data (Cassidy et al., 2018) and may yield different conclusions about the same scientific question (e.g., identifying functional hubs; see Liang et al., 2012). In cases of neuropathology where one might expect distinct gains or losses of functional connections, the use of partial correlation should be interpreted based upon the relative edge density and mean FC. Of the studies reviewed here, 10% used partial correlation, but virtually no study accounted for possible differences in edge density (see Table 2c).

##### Negative edges

Another important aspect of defining edges is how to handle negative FC estimates. Full correlations of RSFC data typically yield an FC distribution in which most edges are positive, but an appreciable fraction are negative. By contrast, partial correlation methods often yield a relative balance between positive and negative FC estimates (e.g., Y. Wang, Kang, Kemmer, & Guo, 2016). There remains little consensus for handling or interpreting negative edge weights in RSFC graph analyses (cf. Murphy & Fox, 2017). In our review, 57% of the studies reported insufficient or no information about how negative edges were handled in graph analyses (Table 2d). Twenty-one percent of studies deleted negative edges prior to analysis, and 9% included the negative weights as positive weights (i.e., using the absolute value of FC). As detailed elsewhere, some graph metrics are either not defined or need to be adapted when negative edges are present (Rubinov & Sporns, 2011).

Importantly, the mean of the marginal association RSFC distribution depends on whether global signal regression (GSR) is included in the preprocessing pipeline (Murphy, Birn, Handwerker, Jones, & Bandettini, 2009). When GSR is included, there is often a balance between positive and negative correlations. If GSR is included as a nuisance regressor, a large fraction of FC estimates may simply be discarded as irrelevant to case-control comparisons, which is a major, untested assumption. The meaning of negative FC, however, remains unclear, with several investigators attributing negative correlations to statistical artifacts and GSR (Murphy et al., 2009; Murphy & Fox, 2017; Saad et al., 2012).

Given that negative correlations are observable in the absence of GSR, however, others have examined whether negative weights contribute differentially to information processing within the network (Parente et al., 2017). Negative correlations may also reflect NMDA action in cortical inhibition (Anticevic et al., 2012). These connections bear consideration given that brain networks composed of only negative connections do not retain a small-world topology, but do have properties distinct from random networks (Parente et al., 2017; Schwarz & McGonigle, 2011). Altogether, the omission of methodological details about negative FC in empirical reports severely hampers the resolution of this important choice point in defining graphs.

#### Degree distribution as a fundamental graph metric

After resolving the questions of node and edge definition, we also believe it is crucial for studies to report information about global network metrics such as characteristic path length, clustering coefficient (aka transitivity), and degree distribution. As noted in Table 3, local and global efficiency were commonly reported (71% and 74%, respectively) and typically across multiple FC thresholds. However, our review revealed that only 27% of studies provided clear descriptive statistics for mean degree, and 16% plotted the degree distribution. In binary graphs, the degree distribution describes the relative frequency of edges for each node in the network. A similar property can be quantified by examining the strength (aka cost) distribution in weighted graphs. We argue that presentation of the degree (strength) distribution in published reports is vital to understanding any RSFC network for a few reasons. First, it provides a sanity check on the data. One current perspective is that human brains are organized to maximize communication while minimizing wiring and metabolic cost (Bassett et al., 2009; Betzel et al., 2017; Chen, Wang, Hilgetag, & Zhou, 2013; Tomasi, Wang, & Volkow, 2013), Thus, when examining whole-brain RSFC data, the most highly connected regions are rare and should be evident in the tail of the degree/strength distribution (Achard, Salvador, Whitcher, Suckling, & Bullmore, 2006). Second, reporting details of the degree distribution facilitates comparisons of graph topology across studies, as well as the impact of preprocessing and analytic decisions such as FC thresholding. Finally, examining the degree distribution as a first step in data analysis may offer otherwise unavailable information about the network topology in healthy and clinical samples. For example, in prior work, we isolated edges from the tail of the degree distribution to understand the impact of the most highly connected, and rare, nodes on the network (Hillary et al., 2014).

**Table 3.** Graph metrics commonly reported in clinical RSFC studies

| **Graph metrics** | ***n* (frequency)** |
| --- | --- |
| Degree Distribution (plotted) | 17 (16.0%) |
| Mean Degree (weighted or binary) | 29 (27.4%) |
| Clustering/Local efficiency | 76 (71.7%) |
| Path Length/Global efficiency | 79 (74.5%) |
| Small worldness | 33(31.1%) |
| Modularity (e.g., Q-value) | 21 (19.8%) |

*Note.* Total frequency is greater than 100% because some studies reported more than one of these metrics.

### Critical Issue 1: Edge Thresholding and Comparing Unequal Networks

We now focus on edge thresholding — that is, how to transform a continuous measure of FC into an edge in the graph. In our review, 39% of studies binarized FC values such that edges were either present or absent in the graph, whereas 45% of studies retained FC as edge weights (see Table 2b). Regardless of whether investigators analyze binary or weighted networks, there are fundamental challenges to comparing networks that differ either in terms of average degree (*k*) or the number of nodes (*N*) (Fornito et al., 2016). In particular, comparing groups on graph metrics such as path length and clustering coefficient can be ambiguous because these metrics have mathematical dependencies on both *k* and *N* (van Wijk et al., 2010).

Most brain parcellation approaches define graphs with an equivalent number of nodes (*N*) in each group. On the other hand, connection density is often a variable of interest in clinical studies where the pathology may alter not only connection strength, but also the number of connections. If *N* is constant, variation in *k* between groups constrains the boundaries of local and global efficiency. If two groups differ systematically in edge density, this almost guarantees between-group differences in metrics such as clustering coefficient and path length. Determining where to intervene in this circular problem has great importance in clinical network neuroscience, where hypotheses often focus on the number and strength of network connections.

To address this issue, several investigators have recommended proportional thresholding (PT) in which the edge density is equated across networks (Achard & Bullmore, 2007; Bassett et al., 2009; Power, Fair, Schlaggar, & Petersen, 2010). Furthermore, to reduce the possibility that findings are not specific to the chosen density threshold, 29% of studies have tested for group differences across a range (e.g., 5-25%; Table 4). However, we argue that defining edges based on PT may not be ideal for clinical studies, where there are often regional differences in functional coupling or pathology-induced alterations in the number of functional connections (Hillary & Grafman, 2017). For example, in depression, the dorsomedial prefrontal cortex exhibits enhanced connectivity with default mode, cognitive control, and affective networks (Sheline, Price, Yan, & Mintun, 2010). As demonstrated below, when FC differs in selected regions between groups, PT is vulnerable to identifying spurious differences in nodal statistics (e.g., degree). The concerns expressed here extend from van den Heuvel and colleagues (2017), who demonstrated that PT increases the likelihood of including spurious connections in the network when groups differ in mean FC.

**Table 4.**
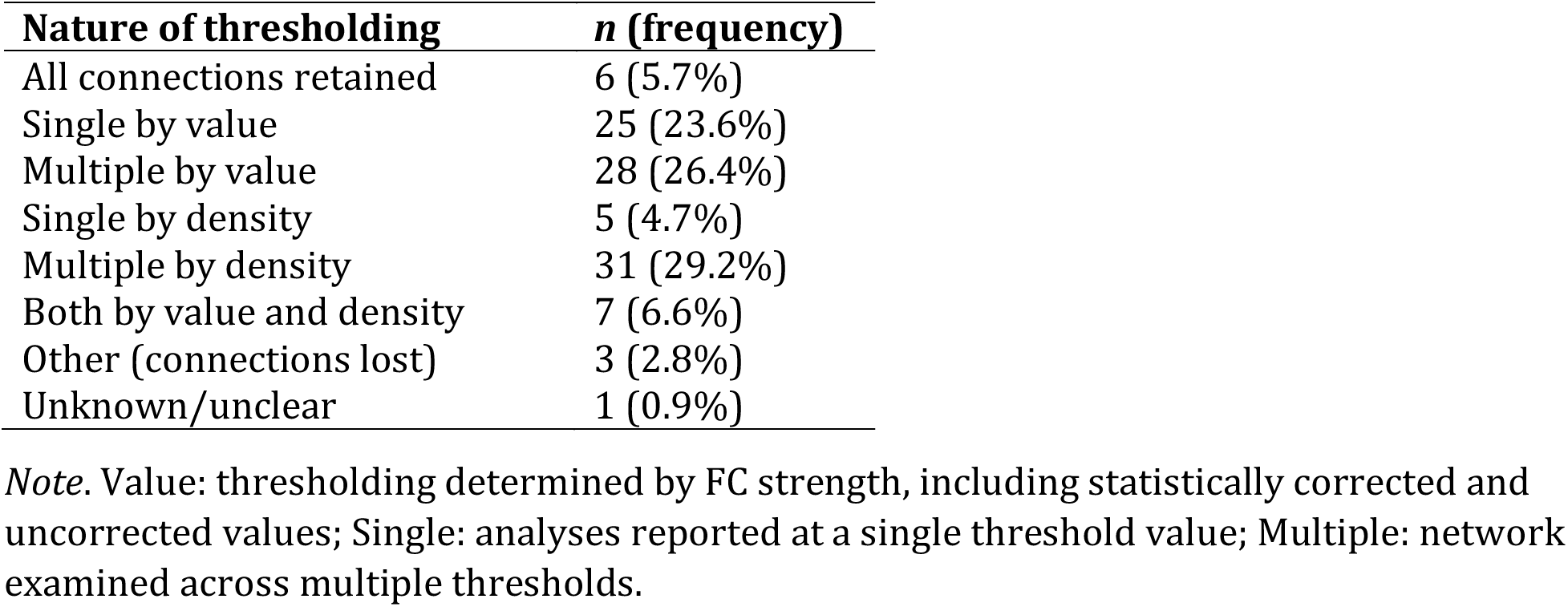
Thresholding method for defining edges in graphs

### Simulation to demonstrate a problem with PT in group comparisons: Whack-a-node

In the first simulation — ‘whack-a-node’ — we examined the consequences of PT on regional inferences when groups differ in FC for selected regions. Unlike empirical resting-state fMRI data, where the underlying causal processes remain relatively unknown, simulations allow one to test the effect of biologically plausible alterations (e.g., hyperconnectivity of certain nodes) on network analyses. Simulations can also clarify the effects of alternative analytic decisions on substantive conclusions in empirical studies. For the details of our simulation approach, see Methods. Briefly, we used a network bootstrapping approach to simulate resting-state network data for a case-control study with 50 patients and 50 controls. We repeated this simulation 100 times, increasing connectivity for patients in three randomly targeted nodes (hereafter called ‘Positive’) and decreasing connectivity slightly, but *nonsignficantly*, in three random nodes (‘Negative’). Our simulations captured both between- and within-person variation in FC changes for targeted nodes (see Methods, Equations 5–7). We compared changes in Positive and Negative nodes to three Comparator nodes that did not differ between groups.

### Results of whack-a-node simulation

#### Proportional thresholding

In a multilevel regression of group difference *t*-statistics (patient – control) on density threshold, we found that PT was sensitive to hyperconnectivity of Positive nodes, reliably detecting group differences, average *t* = 12.4 (*SD* = 1.15), average *p* < .0001 (Figure 2a). Importantly, however, degree was significantly lower in patients than controls for Negative nodes, average *t* = −6.52 (*SD* = 1.15), average *p*< .001. We did not find any systematic difference between groups in Comparator nodes, average *t* = −.22 (*SD* = .65), average *p* = .47.

**Figure 2.**
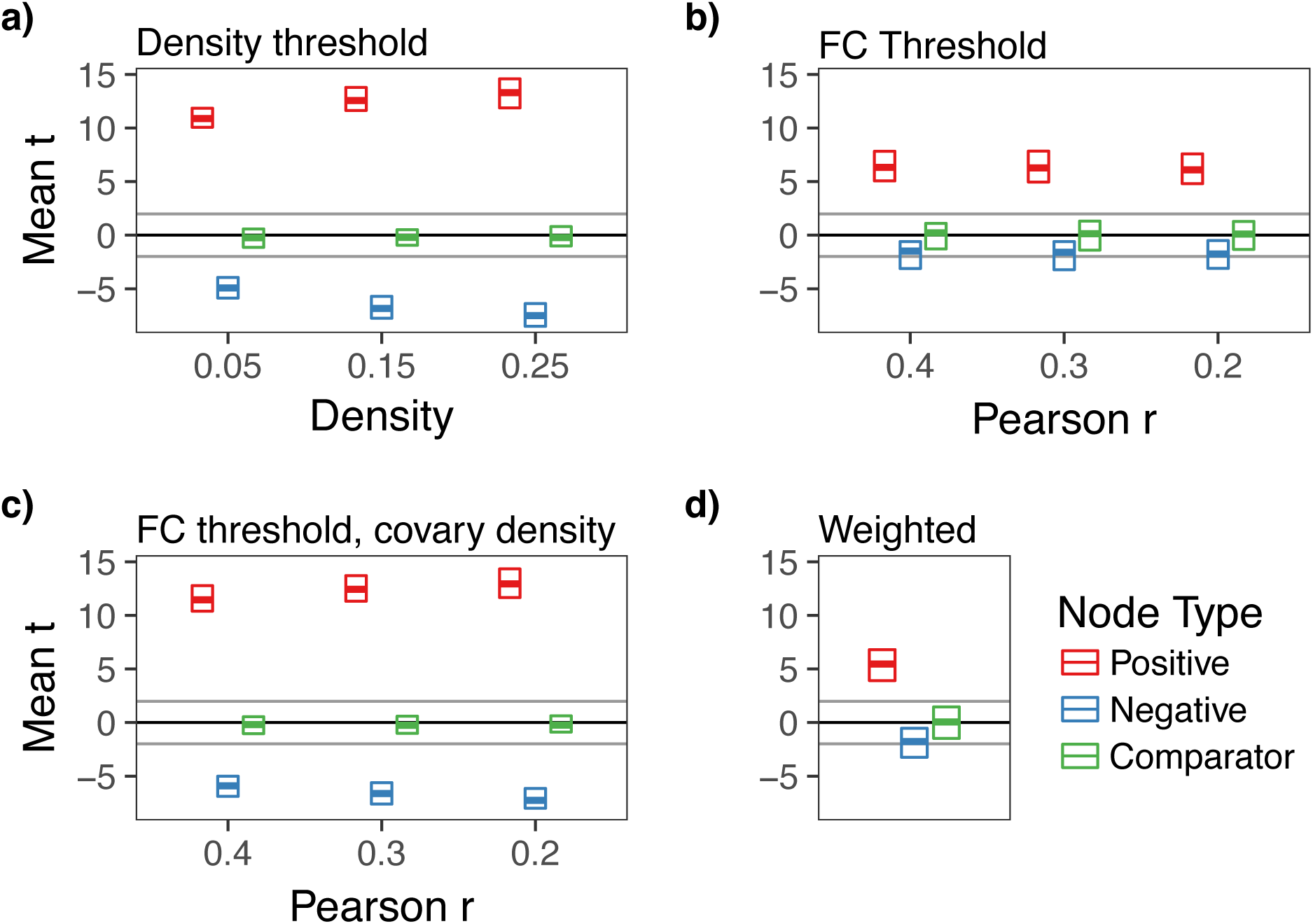
The effect of thresholding method on group differences in degree centrality when there is strong hyperconnectivity in three randomly selected nodes (Positive) and weak hypoconnectivity in three nodes (Negative). Three unaffected nodes are depicted for comparison (Comparator). For parameters of this simulation, see Table S1, Whack-a-node hyperconnectivity. The central bar of each rectangle denotes the median *t* statistic (patient – control) across 100 replication datasets (patient *n* = 50, control *n* = 50), whereas the upper and lower boundaries denote the 90^th^ and 10^th^ percentiles, respectively. The dark line at *t* = 0 reflects no mean difference between groups, whereas the light gray lines at *t* = −1.99 and 1.99 reflect group differences that would be significant at *p* = .05. a) Graphs binarized at 5%, 15%, and 25% density. b) Graphs binarized at *r* = {.2, .3, .4}. c) Graphs binarized at *r* = {.2, .3, .4}, with density included as a between-subjects covariate. d) Strength centrality (weighted graphs).

These group differences were qualified by a significant density x node type interaction (*p* < .0001) such that group differences for Positive nodes were larger at higher densities (Figure 3, top panel), *B* = 11.32 (95% CI = 10.59 – 12.04), *t* = 30.72, *p* < .0001. Conversely, we found equal, but opposite, shifts in Negative nodes (Figure 3, middle panel): group differences became increasingly negative at higher densities, *B* = −11.93, (95% CI = −12.65 − −11.2), *t* = −32.74, *p* < .0001. However, we did not observe an association between density and group differences in Comparator nodes, *B* = .31, *p* = .40 (Figure 3, bottom panel).

**Figure 3.**
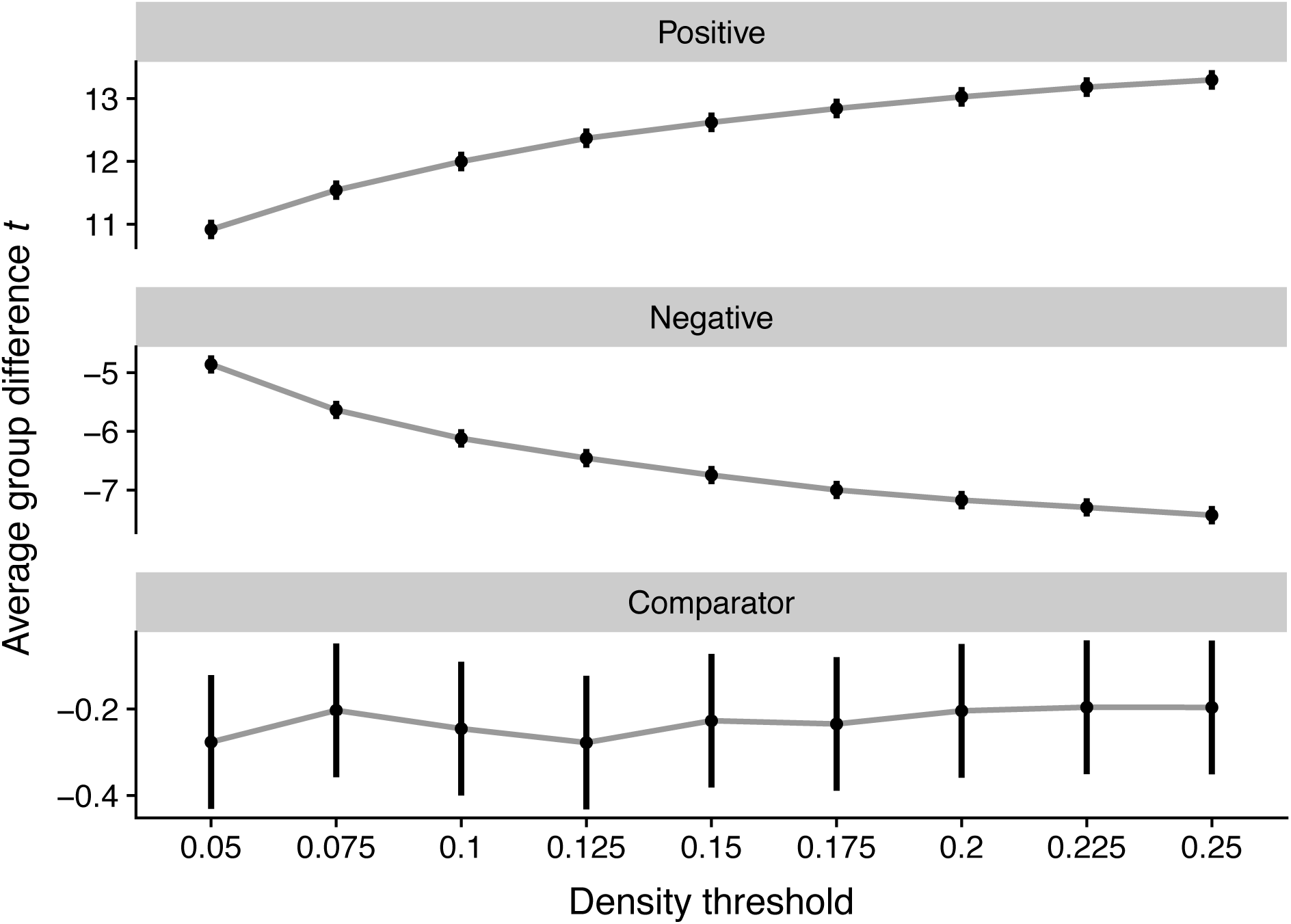
The effect of density threshold on group differences in degree centrality. Dots denote the mean *t* statistic at a given density, whereas vertical lines denote the 95% confidence interval. All statistics reflect group differences in degree centrality computed on graphs binarized at different densities.

#### Functional connectivity (FC) thresholding

In graphs thresholded at differing levels of FC (*r*s ranging between .2 and .5), we found reliable increases in Positive nodes in patients, average *t* = 6.37, average *p* < .0001 (Figure 2b). Negative nodes, however, were not significantly different between groups, average *t* = −1.75, average *p* = .23. Neither did Comparator nodes differ by group, average *t* = .05, *p* =.50. Unlike PT, for FC-thresholded graphs, we did not find a significant threshold x node type interaction, *χ*^2^(16) = 13.38, *p* = .65.

#### Functional connectivity thresholding with density as covariate

By definition, FC thresholding cannot handle the problem of group differences in mean FC. Thus, the risk of FC thresholding alone is that nodal statistics may reflect group differences in mean FC that affect interpretation of topological metrics (e.g., small worldness). To mitigate this concern, one could threshold at a target FC value, then include graph density for each subject as a covariate (as suggested by van den Heuvel et al., 2017). As depicted in Figure 2c, however, although this statistically controls for density, it also reintroduces the constraint that the groups must be equal in average degree. As a result, the pattern of effects mirrors the PT graphs (Figures 2a, 3). More specifically, group differences in Positive nodes were reliably detected, average *t* = 12.39, average *p* < .0001. But patients appeared to be significantly more hypoconnected in Negative nodes compared to controls, average *t* = −6.59, average *p*< .001.

#### Weighted analysis

In analyses of strength centrality, we observed significantly greater degree in Positive nodes, average *t* = 5.5, average *p* = .0003 (Figure 2d). As expected given our simulation design, Negative and Comparator nodes were not significantly different between groups, average *p*s = .19 and .49, respectively.

### Discussion of whack-a-node simulation

In our ‘whack a node’ simulation, three nodes were robustly hyperconnected in ‘patients’, while three nodes were weakly hypoconnected. The principal finding of this simulation was that enforcing equal average degree using PT can spuriously magnify changes in group comparisons of nodal statistics. When the groups were otherwise equal, hyperconnectivity in selected nodes was accurately detected using PT across different graph densities. However, nodes that were weakly hypoconnected tended to be identified as statistically significant. The inclusion of low magnitude, spurious connections into the network inheres from the way in which PT handles the tails of the FC distribution.

By retaining only edges at the high end of the FC distribution, edges that are on the cusp of that criterion are most vulnerable to being removed. For example, at a density of 25%, small variation in FC strength near the 75^th^ percentile could lead to inclusion or omission of an edge. As a result, if FC for edges incident to a given node tends to be weaker in one group than the other, then binary graphs generated using PT will magnify the statistical significance of differences in degree centrality. To the extent that nodal differences in FC strength represent a shift in the central tendency of the distribution, this problem is not solved by applying multiple density thresholds (Figure 3). We observed the same problem if the direction of FC changes was flipped in the simulation: under PT, group comparisons were significant for weakly hyperconnected nodes if some nodes were robustly hypoconnected (see Figure S1). Moreover, our findings were qualitatively unchanged when simulated changes in FC were applied proportionate to the original connection strength, rather than shifting connectivity in correlational units (see Figure S2). This additional simulation respected biologically plausible variation in FC (e.g., greater average connectivity in functional hubs).

We examined FC-based thresholding (here, using Pearson *r* as the metric) and weighted analyses as comparisons to PT. These methods do not suffer from the spurious detection of nodal differences evident under PT. Rather, FC thresholding accurately detected hyperconnected nodes across different thresholds while not magnifying significance of the weakly hypoconnected nodes. However, as noted elsewhere (van den Heuvel et al., 2017; van Wijk et al., 2010), if two groups differ in mean FC, thresholding at a given level (e.g., *r* = .3) in both groups will lead to differences in graph density. This could manifest as widespread differences in nodal statistics due to global differences in the number of edges. We considered whether including per-subject graph density as a covariate in group difference analyses could retain the desirable aspects of FC thresholding while accounting for the possibility of global differences in FC. We found, however, that statistically covarying for density was qualitatively similar in its effects to PT because it constrains the sum of degree changes across the network to be zero between groups (i.e., equal average degree).

The goal of our simulation was to provide a proof of concept that PT may negatively affect nodal statistics in case-control graph studies by enforcing equal average degree. We did not, however, test a range of parameters to identify the conditions under which this concern holds true. The simulation focused specifically on degree centrality in the binary case and strength in the weighted case. While untested, we anticipate that these effects likely generalize to other nodal measures such as eigenvector centrality. Importantly, the problems with PT highlighted above occur regardless of edge density (Figure 3), so the use of multiple edge densities does not adequately address the “whack-a-node” issue.

### Critical Issue 2: Matching theory to scale: Telescoping levels of analysis in graphs

The second major methodological theme from our literature review concerns the alignment between neurobiological hypotheses and graph analyses. We refer to this as theory-to-scale matching. RSFC graphs offer telescoping levels of information about intrinsic connectivity patterns, from global information such as average path length to details such as connectivity differences in a specific edge. For example, in major depression, resting-state studies have focused analyses on specific subnetworks composed of the dorsal medial prefrontal cortex, anterior cingulate cortex (ACC), amygdala, and medial thalamus (for a review, see L. Wang, Hermens, Hickie, & Lagopoulos, 2012).

We propose that graph analyses should be conceptualized and reported in terms of telescoping levels of analysis from global to specific: global topology, modular structure, nodal effects, edge effects. Our review of the clinical network neuroscience literature revealed that the majority of studies (69%; see Table 5) tested whether groups differed on global metrics such as small-worldness. Importantly, however, many studies provided limited theoretical justification for why the pathophysiology of a given brain disorder should alter the global topology of the network. In the following, we provide more specific comments on path length and small-worldness, and how investigators might ideally match hypotheses to levels of analysis.

**Table 5.** Level of group difference hypotheses in graph analyses (i.e., telescoping)

| **Hypothesis level** | ***n* (frequency)** |
| --- | --- |
| Global | 73 (68.9%) |
| Module (subnetwork) | 41 (38.7%) |
| Nodal (Region) | 52 (49.1%) |
| Edge | 1 (0.9%) |

*Note.* Global: examining whole-graph network features (e.g., small-worldness), Module: examining subnetworks/modules (e.g., default mode network). Total frequency is greater than 100% because some studies tested hypotheses at multiple levels.

### The clinical meaningfulness of small-worldness

In a defining study for network neuroscience,Watts and Strogatz (1998) demonstrated that the organization of the central nervous system in *C. elegans* reflected a “small-world” topology (cf. Muldoon, Bridgeford, & Bassett, 2016). The impact of this finding continues to resonate in the network neuroscience literature 20 years later (Hilgetag & Kaiser, 2004; Sporns & Zwi, 2004), with many studies focusing on “disconnection” and the loss of network efficiency as quantified by small-worldness (31% of the studies reviewed here). Although small-world topologies have been observed in most studies of brain function (Bassett & Bullmore, 2016), the relevance of this organization for facilitating human information processing remains unclear. Other features of human neural networks, such as modularity, may have more important implications for network functioning (Hilgetag & Goulas, 2015). Higher network modularity reflects a graph in which the connections among nodes tend to form more densely connected communities, which tend to be robust to random network disruption (Shekhtman, Shai, & Havlin, 2015).

It remains uncertain that, as a general rule, brain pathology *should* be reflected in a measure of small-worldness. For example, a small-world topology is preserved even in experiments that dramatically reduce sensory processing via anesthesia in primates (Vincent et al., 2007) and in disorders of consciousness in humans (Crone et al., 2014). Recognizing that conventional measures of small-worldness (e.g.,Humphries & Gurney, 2008) depend on density and do not handle variation in connection strength, recent research has recast this concept and its quantification in terms of the “small world propensity” of a network (Muldoon et al., 2016).

Although we do not dispute the value of global graph metrics such as small-worldness as important descriptors of a network organization, by definition, they provide information at only the most macroscopic level. Group differences in global metrics may largely reflect more specific effects at finer levels of the graph. For example, removing connections in functional hub regions selectively tends to reduce global efficiency and clustering (Hwang, Hallquist, & Luna, 2013). Likewise, failing to identify group differences in global structure does not imply equivalence at other levels of the graph (e.g., nodes or modules). To demonstrate the point that substantial group differences in finer levels of the graph may not be evident in global metrics, we conducted a simulation in which the groups differed substantially in modular connectivity.

### Simulation to demonstrate the importance of understanding graphs at multiple levels: global insensitivity to modular effects

Extending the basic approach of our whack-a-node simulation, we used a 13-module parcellation of the 264-node groundtruth graph in a simulation of 50 ‘controls’ and 50 ‘patients’ (modular structure from Power et al., 2011). More specifically, we increased FC in the fronto-parietal network (FPN) and dorsal attention network (DAN) in controls, and increased FC in the default mode network (DMN) in patients. The simulation primarily examined group differences in small-worldness (global metric) and within-and between-module degree (nodal metrics). Additional details are provided in the Methods.

#### Results of global insensitivity simulation

Consistent with common methods in the field, we applied proportional thresholding (PT) between 7.5% and 25% to each graph. We computed small-worldness, *σ*, at each threshold (see Methods), then tested for group differences in this coefficient. As depicted in Figure 4, we did not observe any significant group differences in small-worldness at any edge density, *M t* = .36, *p*s > .3.

**Figure 4.**
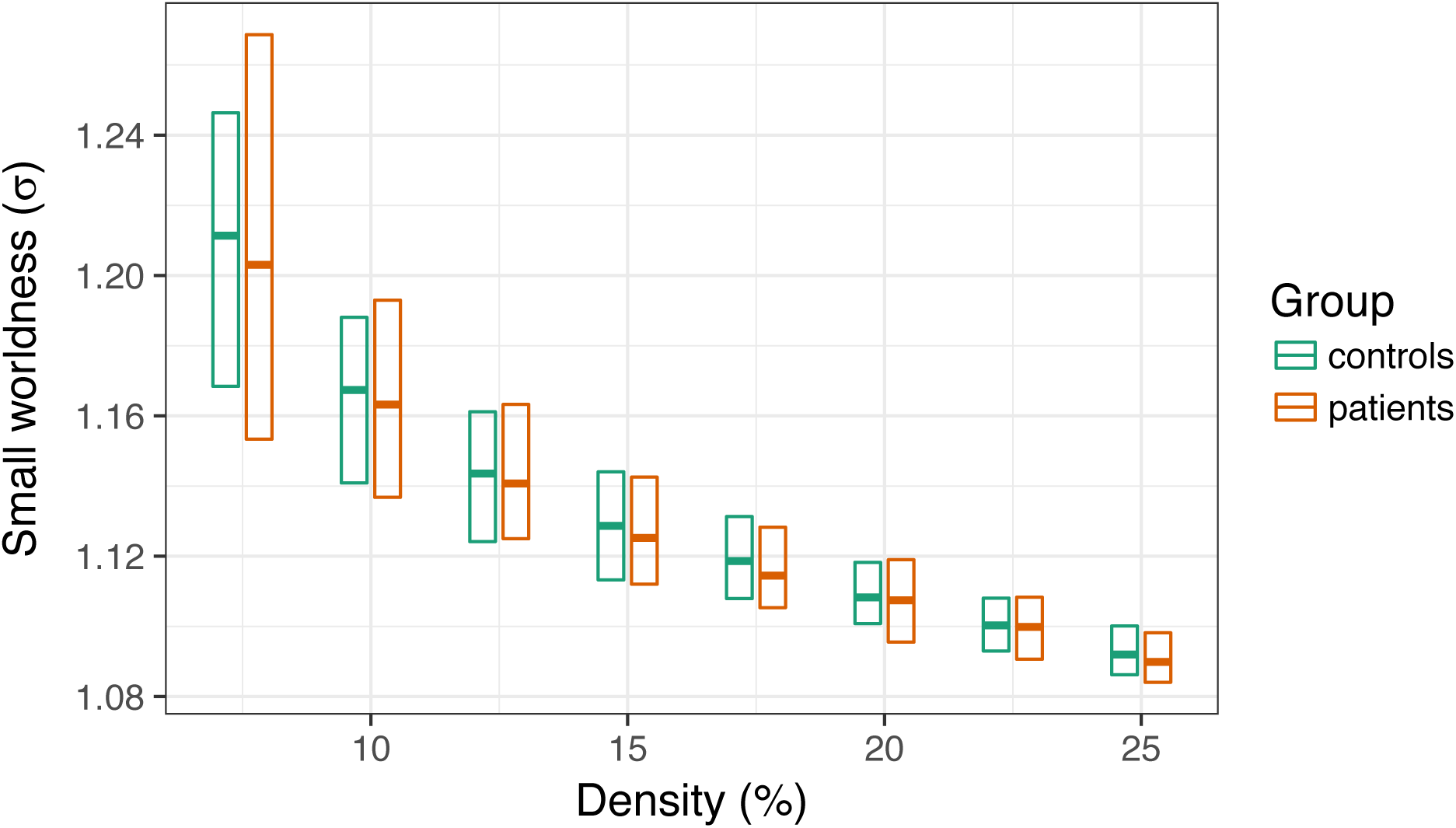
Group differences in small worldness (σ) as a function of edge density. The central bar of each rectangle denotes the median σ statistic (patient – control), whereas the upper and lower boundaries denote the 90^th^ and 10^th^ percentiles, respectively.

However, consistent with the structure of our simulations, we found large group differences in within- and between-module degree (Figure 5). In a multilevel regression of within-network degree on group, density, and module, we found a significant DMN increase in patients irrespective of density, *B* = 1.37, *t* = 33.25, *p* < .0001. Likewise, controls had significantly higher within-network degree in the FPN and DAN, *t*s = 21.67 and 13.52, respectively, *p*s < .0001. These findings were mirrored in group analyses of between-network degree in the DMN, FPN, and DAN, *p*s < .0001 (see Figure 5).

**Figure 5.**
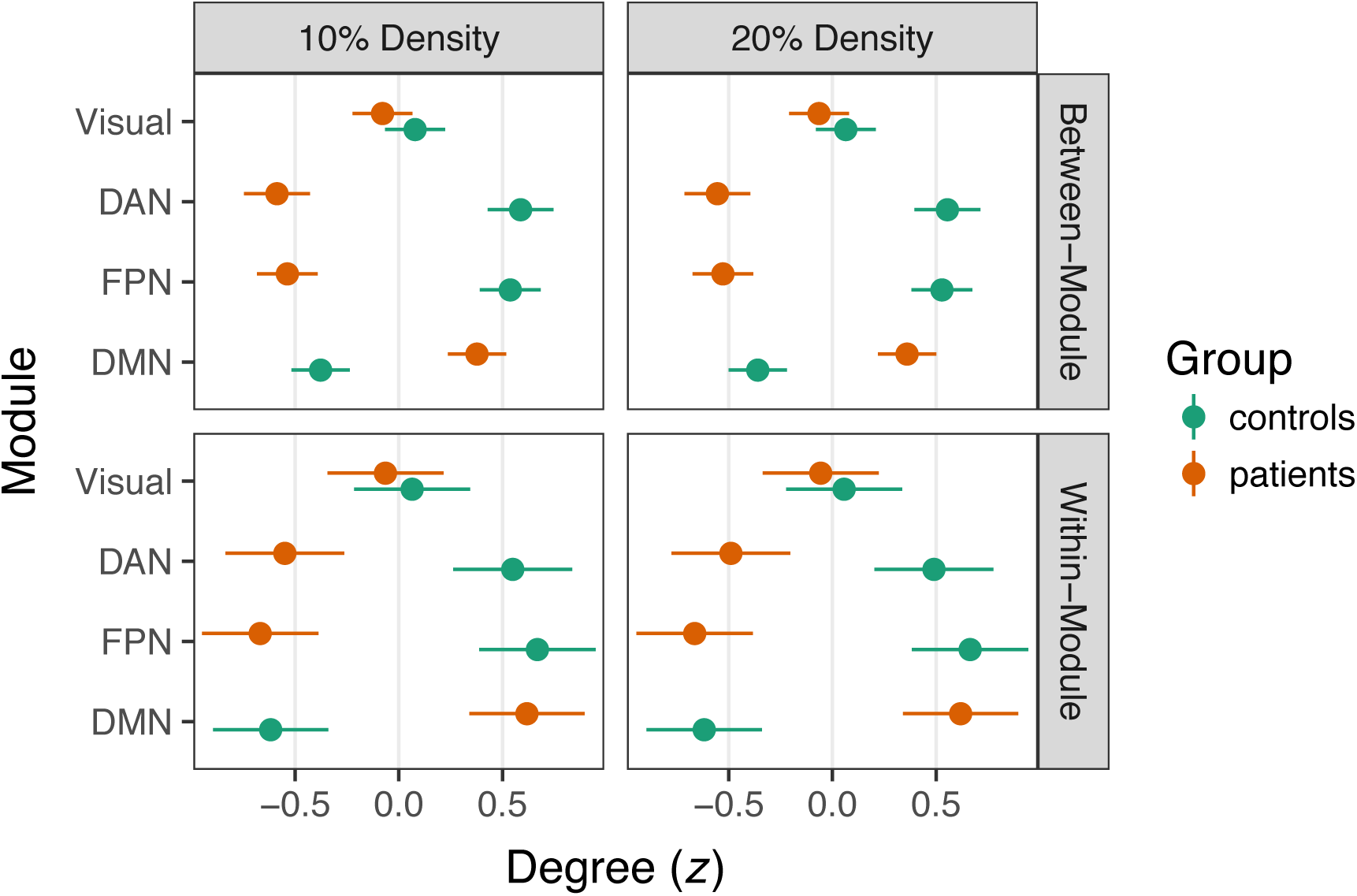
Group differences in *z*-scored degree statistics at 10 and 20% density. Degree differences for connections between modules are depicted in the top row, whereas within-module differences are in the bottom row. Dots denote the mean *z* statistic across nodes, whereas the lines represent the 95% confidence interval around the mean. Nonsignificant group differences in the Visual network are depicted for comparison, whereas connectivity in the DAN, FPN, and DMN was focally manipulated in the simulation. DAN = Dorsal Attention Network; FPN = Fronto-Parietal Network; DMN = Default Mode Network.

#### Discussion of global insensitivity simulation

In the global insensitivity simulation, we induced large group changes in FC at the level of functional modules that represent canonical resting-state networks (e.g., the DMN). The simulation differentially modulated FC within and between regions of the DAN, FPN, and DMN. In group analyses of within- and between-module degree centrality, we detected these large shifts in FC. However, despite robust differences in network structure, the two groups were very similar in the small-world properties of their graphs.

As with the whack-a-node simulation, we did not seek to test the range of conditions under which these findings would hold. Rather, the global insensitivity simulation provides a proof of concept that researchers should be aware that the absence of group differences at a higher level of the graph (here, global topology) does not suggest that the networks are otherwise similar at lower levels (here, modular connectivity). As we have noted above, in graph analyses of case-control resting-state networks, we encourage researchers to state their study goals in terms that clearly match the hypotheses to the scale of the graph.

In the global insensitivity simulation, failing to detect differences in small-worldness should not be seen as an omnibus test of modular or nodal structure. Likewise, if group differences are detected at the global level, there may be substantial value in interrogating finer differences in the networks, even if these were not hypothesized a priori.

### General Summary

The overarching goal of this review was to promote shared standards for reporting findings in clinical network neuroscience. Our survey of the resting-state functional connectivity literature revealed the popularity and promise of graph theory approaches to network organization in brain disorders. This potential is evident in large-scale initiatives for acquiring and sharing resting-state data in different populations (e.g., the Human Connectome Project; Barch et al., 2013). Publicly available resting-state data can also support reproducibility efforts by serving as replication datasets to corroborate specific findings in an independent sample (e.g.,Jalbrzikowski et al., 2017). We share the field’s enthusiasm for such work and anticipate that with further methodological refinement and standardization, the coupling of network science and brain imaging can provide novel insights into the neurobiological basis of brain disorders.

Our review, however, suggested that the heterogeneity of methods is preventing the field from realizing its potential. Graph analyses across clinical studies varied substantially in terms of brain parcellation, FC quantification, and the use of thresholding methods to define edges in graphs. These decisions are fundamental to graph theory and precede analyses at specific levels such as global topology. In addition, although it was not a focus of our review, there was substantial variation in what network metrics were reported across studies. The lack of standardization in methods at multiple decision points has multiplicative consequences: the likelihood that any two studies used the same parcellation scheme, FC definition, thresholding strategy, and network metric was remarkably low. This makes formal meta-analyses of the clinical network neuroscience literature virtually impossible at the present time, detracting from efforts to distinguish distinct pathophysiological mechanisms or to identify transdiagnostic commonalities. In addition, methodological heterogeneity in graph analyses undercuts the value of data sharing efforts that have made massive datasets available to the network neuroscience community. For the immense potential of data sharing to be realized, standardization must occur not only in data acquisition, but also in data analysis, with a shared framework to guide hypothesis-to-scale matching in graphs.

As summarized in Box 2, we believe that the field should work toward a principled, common approach to graph analyses of RSFC data. This is a challenging proposition because of the rapid and exciting developments in functional brain parcellation (e.g.,Schaefer et al., in press), FC definition (e.g.,Cassidy, Rae, & Solo, 2015), edge thresholding (van den Heuvel et al., 2017), and network metrics (e.g.,Vargas & Wahl, 2014). Such developments highlight both the enthusiasm for, and relative infancy of, network neuroscience as a field. Although we are sensitive to the importance of continuing to refine functional parcellations of the brain, we also see great value in developing field-standard parcellations to promote comparability. Indeed, many aspects of network structure (e.g., homogeneity of functional connectivity patterns within a region) are largely convergent above a certain level of detail (likely 200-400 nodes) in the functional parcellation (Craddock et al., 2012; Schaefer et al., in press). Likewise, the optimal approach for quantifying functional connectivity is an open question (Smith et al., 2011), yet in the absence of methodological convergence, graphs were often not comparable across studies.

A related dilemma was that in 57% of studies, little or no detail was provided about how negative FC values were incorporated into graph analyses. This was especially troubling insofar as global signal regression tends to yield an FC distribution in which approximately half of edges are negative (Murphy et al., 2009). Furthermore, FC estimates at the low end of the distribution may have different topological properties such as reduced modularity (Schwarz & McGonigle, 2011). Even among the 21% of studies in which negative edges were explicitly dropped, it remains unclear what consequences this decision has on substantive conclusions about graph structure. In recent years, there have been advances in quantifying common graph metrics such as modularity in weighted networks that include negative edges (Rubinov & Sporns, 2011), as well as increasing calls for weighted, not binary, graph analyses (Bassett & Bullmore, 2016). Regardless, we believe that greater clarity in reporting of negative FC will promote comparisons among clinical studies.

Methodological heterogeneity also resulted in very few graph statistics that were reported in common across studies, an essential ingredient for examining the reproducibility of findings. Networks that are resilient to the deletion of specific edges often have a highly skewed degree distribution (Callaway, Newman, Strogatz, & Watts, 2000) that may relate to small-world network properties (Achard et al., 2006). We propose that studies should routinely depict this distribution. Likewise, broad metrics such as edge density, mean FC, transitivity, and characteristic path length provide important information about the basic properties of graphs that contextualize more detailed inferential analyses. The challenge of developing reporting standards in clinical network neuroscience echoes the broader conversation in neuroimaging about reproducibility, especially the importance of detail and transparency in the analytic approach (Nichols et al., 2017).

In addition to the general issues of standardizing graph analyses and reporting procedures, our review examined two critical issues in greater detail. First, we considered the potential benefits and risks of using proportional thresholding (PT), a common procedure for equating the number of edges between graphs. Second, we articulated the value of considering the telescoping levels of graph structures in order to match a hypothesis to the corresponding scale of the graph.

Roughly one third of the studies included in our review applied PT, thresholding graphs at single or multiple edge densities. Although this approach is aligned with prior work highlighting concerns about comparing unequal networks (van Wijk et al., 2010), its application in clinical studies is often conceptually problematic. Many brain pathologies appear to affect targeted regions or networks, while leaving the connectivity of other regions largely undisturbed. For example, although frontolimbic circuitry is heavily implicated in mood disorders (Price & Drevets, 2010), visual networks largely appear unaffected. There is growing evidence that brain disorders alter the strength of functional coupling and potentially the number of functional connections (Hillary & Grafman, 2017). Consequently, if some regions are affected by the pathology, but others are similar to a matched control population, PT may erroneously remove or add connections to graphs in one group in order to maintain equal average degree between groups. Furthermore, if a brain pathology alters the density of functional connections — for example, neurological disruption is associated with hyperconnectivity (Hillary et al., 2015) — PT will preclude the investigator detecting density differences between groups. If, in truth, the groups differ in edge density, artificially equating density also detracts from the interpretability of graph analyses (cf. van den Heuvel et al., 2017).

In addition to these conceptual problems, our whack-a-node simulation demonstrated that PT may result in the detection of spurious group differences (Figure 2a). Altogether, applying PT in clinical studies may be a methodological double jeopardy, characterized by reduced sensitivity to pathology-related differences in connection density and the risk of identifying nodal differences between groups that are a statistical artifact. These risks make PT especially unappealing when one considers that FC-based thresholding and weighted analyses accurately detected group differences (Figure 2b,d) while also allowing edge density to vary. We acknowledge, however, that a previous empirical study found that group differences were more consistent across multiple thresholds using proportional, as opposed to FC-based, thresholding (Garrison, Scheinost, Finn, Shen, & Constable, 2015).

We therefore have two recommendations for edge thresholding in case-control comparisons. First, weighted analyses or FC thresholding should typically be preferred to PT if one is interested in nodal statistics. Second, to rule out the possibility that nodal findings reflect global differences in mean FC, one could include mean FC as a covariate in weighted analyses or per-subject density in analyses of FC-thresholded binary graphs. Crucially, we propose that these be treated as sensitivity analyses conducted only after establishing a nodal group difference. That is, if one identifies group differences in FC-thresholded graphs (e.g., greater degree in anterior cingulate cortex among patients), does including edge density as a covariate abolish this finding? If so, it suggests that differences in global topology may account for the nodal finding. However, one should not include density as a covariate in FC-thresholded graphs as a first step to identify which nodes differ between groups, as this could fall prey to the ‘whack a node’ problem (i.e., spurious nodal effects).

The second critical issue was that many studies provided limited theoretical justification for the alignment between a given hypothesis and the corresponding graph analysis. A majority of studies (69%) tested whether groups differed in global metrics such as small-worldness, but most pathologies (e.g., brain injury, Alzheimer’s disease) primarily affect regional hubs within networks (Crossley et al., 2014). Our global insensitivity simulation focused on the importance of matching hypotheses to graph analyses, or telescoping. We demonstrated that global graph metrics, specifically small-worldness, may not be sensitive to group differences in module or node centrality. Metrics such as modularity and nodal centrality offer vital information about regional brain organization that can be interpreted in the context of alterations in average degree. The general point is that one cannot generalize findings from one level of a graph to another, nor should null effects at one level be viewed as suggesting that the groups are similar at other levels. By conceptualizing a data analysis plan in terms of the telescoping structure of graphs, researchers can clearly delineate confirmatory from exploratory analyses, which is consistent with the spirit of reproducible science in neuroimaging (Poldrack et al., 2017).

##### Box 2: Recommendations for Best Practices and Convergence in Clinical Network Neuroscience

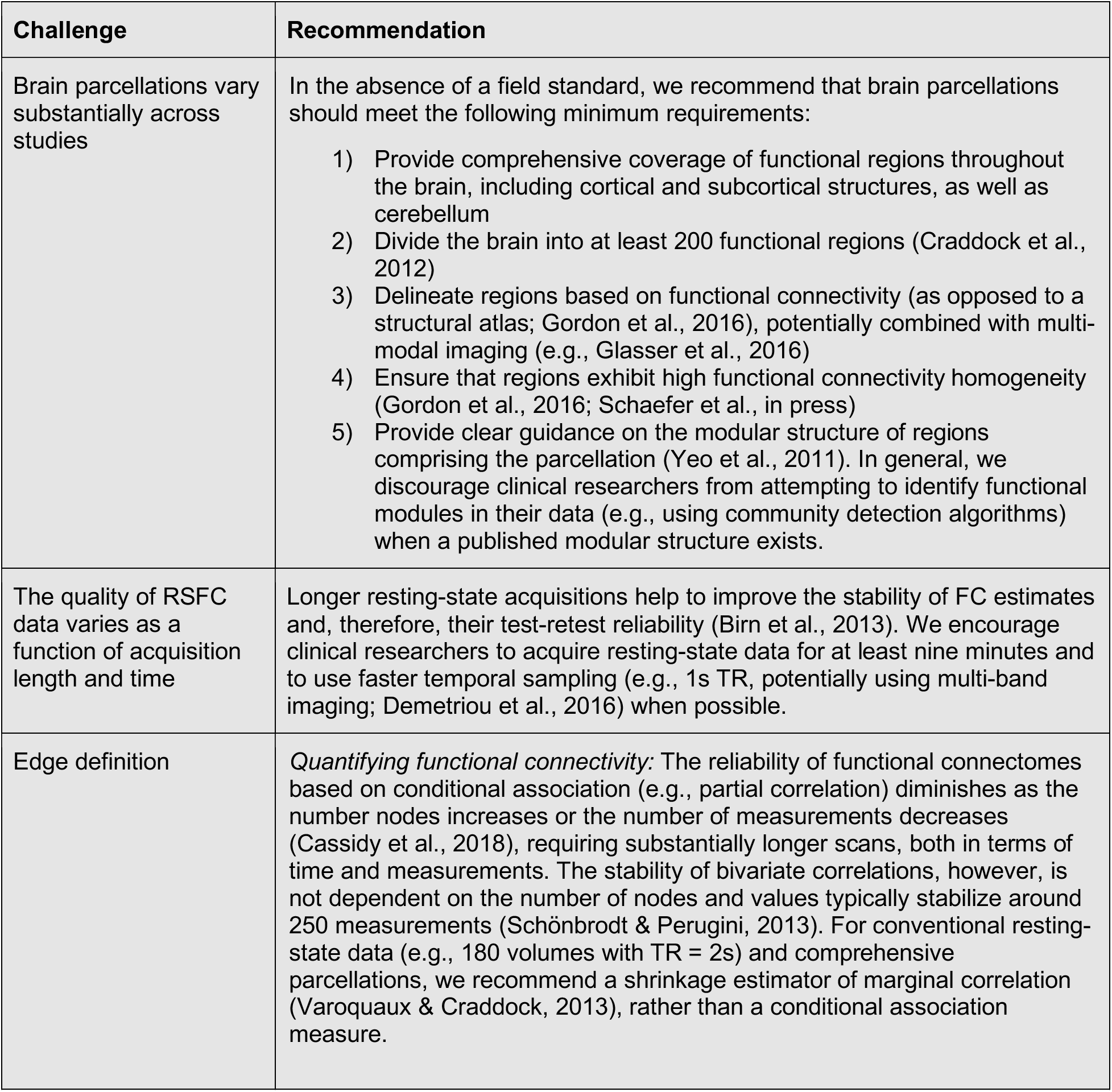
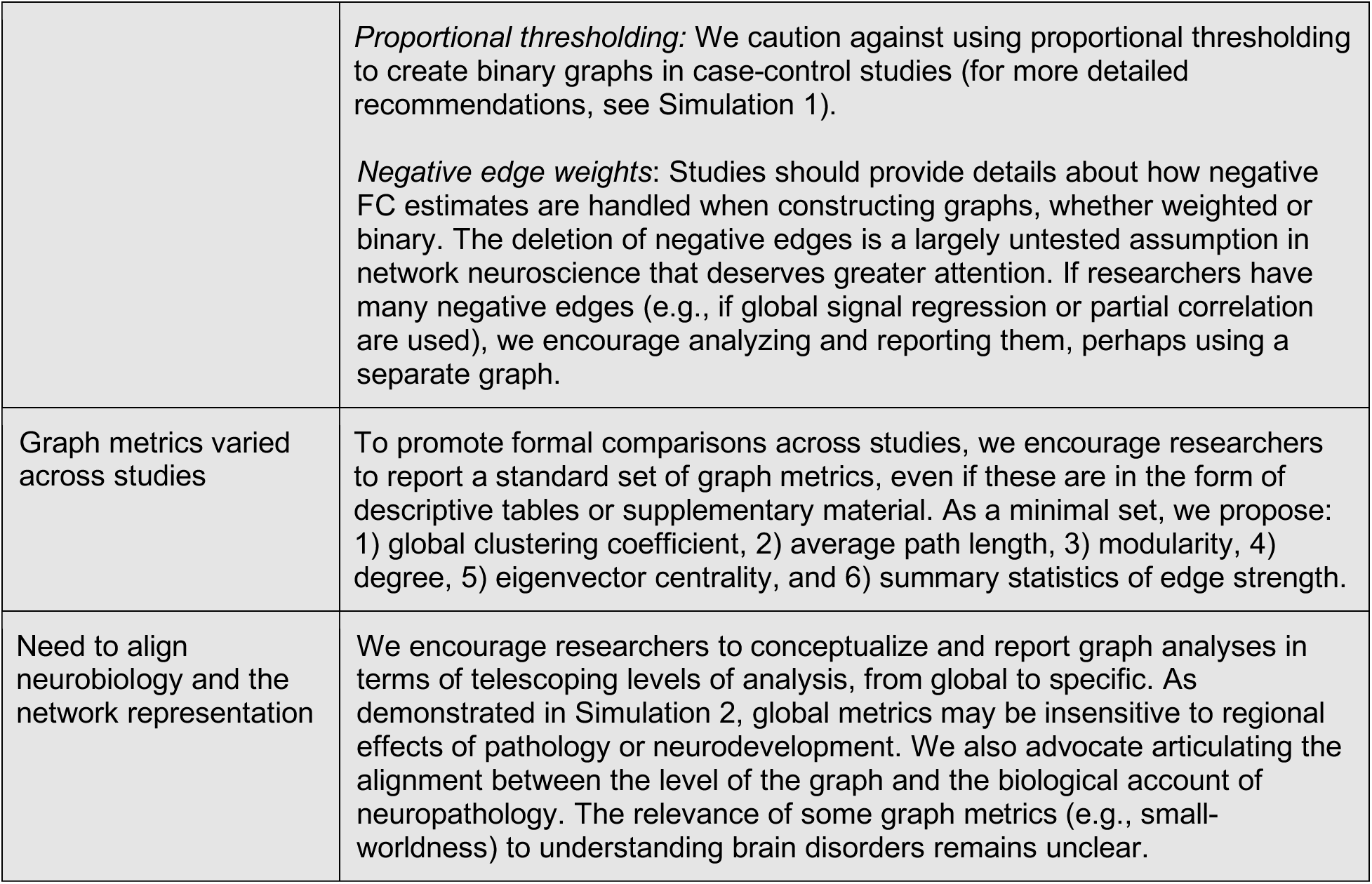

Neuroscience
| Challenge | Recommendation |
| --- | --- |
| Brain parcellations vary substantially across studies | In the absence of a field standard, we recommend that brain parcellations should meet the following minimum requirements:<br>1) Provide comprehensive coverage of functional regions throughout the brain, including cortical and subcortical structures, as well as cerebellum<br>2) Divide the brain into at least 200 functional regions (Craddock et al., 2012)<br>3) Delineate regions based on functional connectivity (as opposed to a structural atlas; Gordon et al., 2016), potentially combined with multi-modal imaging (e.g., Glasser et al., 2016)<br>4) Ensure that regions exhibit high functional connectivity homogeneity (Gordon et al., 2016; Schaefer et al., in press)<br>5) Provide clear guidance on the modular structure of regions comprising the parcellation (Yeo et al., 2011). In general, we discourage clinical researchers from attempting to identify functional modules in their data (e.g., using community detection algorithms) when a published modular structure exists. |
| The quality of RSFC data varies as a function of acquisition length and time | Longer resting-state acquisitions help to improve the stability of FC estimates and, therefore, their test-retest reliability (Birn et al., 2013). We encourage clinical researchers to acquire resting-state data for at least nine minutes and to use faster temporal sampling (e.g., 1s TR, potentially using multi-band imaging; Demetriou et al., 2016) when possible. |
| Edge definition | *Quantifying functional connectivity:* The reliability of functional connectomes based on conditional association (e.g., partial correlation) diminishes as the number nodes increases or the number of measurements decreases (Cassidy et al., 2018), requiring substantially longer scans, both in terms of time and measurements. The stability of bivariate correlations, however, is not dependent on the number of nodes and values typically stabilize around 250 measurements (Schönbrodt & Perugini, 2013). For conventional resting-state data (e.g., 180 volumes with TR = 2s) and comprehensive parcellations, we recommend a shrinkage estimator of marginal correlation (Varoquaux & Craddock, 2013), rather than a conditional association measure. |
|  | *Proportional thresholding:* We caution against using proportional thresholding to create binary graphs in case-control studies (for more detailed recommendations, see Simulation 1).<br>*Negative edge weights:* Studies should provide details about how negative FC estimates are handled when constructing graphs, whether weighted or binary. The deletion of negative edges is a largely untested assumption in network neuroscience that deserves greater attention. If researchers have many negative edges (e.g., if global signal regression or partial correlation are used), we encourage analyzing and reporting them, perhaps using a separate graph. |
| Graph metrics varied across studies | To promote formal comparisons across studies, we encourage researchers to report a standard set of graph metrics, even if these are in the form of descriptive tables or supplementary material. As a minimal set, we propose: 1) global clustering coefficient, 2) average path length, 3) modularity, 4) degree, 5) eigenvector centrality, and 6) summary statistics of edge strength. |
| Need to align neurobiology and the network representation | We encourage researchers to conceptualize and report graph analyses in terms of telescoping levels of analysis, from global to specific. As demonstrated in Simulation 2, global metrics may be insensitive to regional effects of pathology or neurodevelopment. We also advocate articulating the alignment between the level of the graph and the biological account of neuropathology. The relevance of some graph metrics (e.g., small-worldness) to understanding brain disorders remains unclear. |

In summary, there is a need for a common framework to inform graph theory analyses of RSFC data in the clinical neurosciences. Any recommendations should emerge organically from a scientific community whose investigators voluntarily adopt procedures that maximize sensitivity to hypothesized effects and simultaneously permit graph comparisons among studies (cf. the COBIDAS effort in neuroimaging more broadly; Nichols et al., 2017). The implementation of best practices in task-based fMRI (e.g., handling autocorrelation; Woolrich, Ripley, Brady, & Smith, 2001) has been supported by powerful, usable, and free software. Although similar software is increasingly available for resting-state studies (e.g.,Chao-Gan & Zang Yu-Feng, 2010; Whitfield-Gabrieli & Nieto-Castanon, 2012), best practices are still emerging, and we believe that software developers will be crucial to promoting a standardized graph theory analysis pipeline. We anticipate that a common methodological framework will promote hypothesis-driven research, alignment between theory and graph analysis, reproducibility, data sharing, meta-analyses, and ultimately more rapid progress of clinical network neuroscience.

## Methods

### Pubmed Search Syntax

#### Neurological disorders query

(graph OR graphical OR graph-theor* OR topology) AND (brain OR fMRI OR connectivity OR intrinsic) AND (resting-state OR resting OR rest) AND (neurological OR brain injury OR multiple sclerosis OR epilepsy OR stroke OR CVA OR aneurysm OR Parkinson’s OR MCI or Alzheimer’s OR dementia OR HIV OR SCI OR spinal cord OR autism OR ADHD OR intellectual disability OR Down syndrome OR Tourette) AND “humans”[MeSH Terms]

#### Mental disorders query

(graph OR graphical OR graph-theor* OR topology) AND (brain OR fmri OR connectivity OR intrinsic) AND (resting-state OR resting OR rest) AND (clinical OR psychopathology OR mental disorder OR psychiatric OR neuropsychiatric OR depression OR mood OR anxiety OR addiction OR psychosis OR bipolar OR borderline OR autism) AND “humans”[MeSH Terms]

### General Approach to Network Simulations

To approximate the structure of functional brain networks, we identified a young adult female subject who completed a 5-minute resting-state scan in a Siemens 3T Trio Scanner (TR = 1.5s, TE=29ms, 3.1 × 3.1 × 4.0mm voxels) with essentially no head movement (mean framewise displacement [FD] = .08mm; max FD = .17mm). We preprocessed the data using a conventional pipeline, including: 1) motion correction (FSL mcflirt); 2) slice timing correction (FSL slicetimer); 3) nonlinear deformation to the MNI template using the concatenation of functional → structural (FSL flirt) and structural → MNI152 (FSL flirt + fnirt) transformations; 4) spatial smoothing with a 6mm FWHM filter (FSL susan); 5) and voxelwise intensity normalization to a mean of 100. After these steps, we also simultaneously applied nuisance regression and bandpass filtering, where the regressors were six motion parameters, average CSF, average WM, and the derivatives of these (16 total regressors). The spectral filter retained fluctuations between .009 and .08 Hz (AFNI 3dBandpass). We then estimated functional connectivity (FC) of 264 functional ROIs from the Power and colleagues parcellation, where each region was defined by a 5mm radius sphere centered on a specific coordinate. The activity of a region over time was estimated by the first principal component of voxels in each ROI, and the functional connectivity matrix (264 × 264) was estimated by the pairwise Pearson correlations of time series.

This 264 × 264 adjacency matrix, **W**, served as the groundtruth for all simulations, with specific within-person, between-person, and between-group alterations applied according to a multilevel simulation of variation across individuals (details of simulation parameters provided in Table S1). More specifically, to approximate population-level variability in a case-control design, we simulated resting-state adjacency matrices for 50 ‘patients’ and 50 ‘controls’ by introducing systematic and unsystematic sources of variability for each simulated participant. Systematic sources were intended to test substantive hypotheses about proportional thresholding (whack-a-node simulation) and insensitivity to global versus modular differences (global insensitivity simulation), whereas unsystematic sources reflected within- and between-person variation in edge strength.

In this model, the simulated edge strength between two nodes, *i* and *j*, for a given subject, *s*, is:

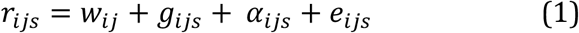

where *w*_*ij*_ is the edge strength from the groundtruth adjacency matrix **W**. Global variation in mean FC is represented by *g*_*ijs*_, which reflects contributions of both between- and within-person variation:

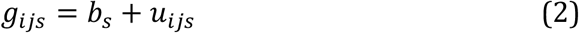

where *b*_*s*_ represents normally distributed between-person variation in mean FC:

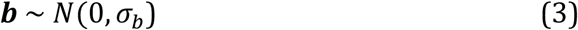

while *σ_b_* controls the level of between-person variation in the sample. The term *u*_*ijs*_ represents within-person variation of this edge relative to the person mean FC, *b*_*s*_. Within-person variation in FC across all edges is assumed to be normally distributed:

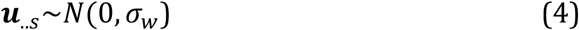

with *σ*_*w*_ scaling the degree of within-person FC variation across all edges.

Node-specific shifts in FC are represented by *α*_*ijs*_, which includes both between-person and within-node components. More specifically, the modulation of FC between nodes *i* and *j* is given by:

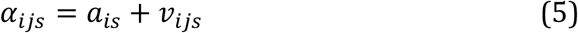

where between-person variation in FC for node *i* is:

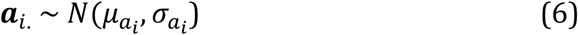

with *μ*_*a*_*i*__ and *σ*_*a*_*i*__ capturing the mean and standard deviation in FC shifts for node *i* across subjects, respectively. Edgewise FC variation of a node *i* across its neighbors, *j*, is given by:

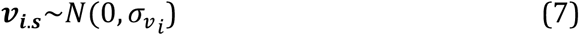

where *σ*_*v*_*i*__ represents the standard deviation of FC shifts across neighbors of *i*. When nodes *i* and *j* were both manipulated, the shifts were applied sequentially such that FC for the edge between *i* and *j* was not allowed to have compounding changes. That is, we set *α*_*ijs*_ = 0 for *i* > *j*.

Finally, *e*_*ijs*_ represents the random variation in FC for the edge between *i* and *j* for subject *s*. This variation was assumed to be normally distributed across all edges for a subject:

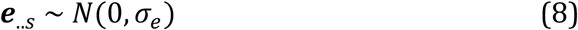

where *σ*_*e*_ controls the standard deviation of edge noise across subjects.

### Whack-a-Note Simulation Methods

As mentioned above, proportional thresholding (PT) is often applied in case-control studies to rule out the possibility that network differences between groups reflect differences in the total number of edges. Importantly, in graphs of equal order *N* (i.e., the same number of nodes), proportional thresholding equates both the density, *D*, and average degree, 〈*k*〉 between subjects:

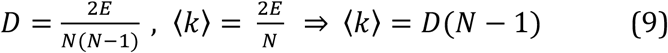

where *E* denotes the number of unique edges in an undirected graph with no self-loops. As noted by van den Heuvel and colleagues (2017), when one applies PT, differences in average connectivity strength can lead to the inclusion of weaker edges in more sparsely connected groups. Furthermore, weak edges estimated by correlation are more likely to reflect an unreliable relationship between nodes. Thus, if one group has lower mean functional connectivity, PT could introduce spurious connections, potentially undermining group comparisons of network topology.

That is, when groups are otherwise equivalent, the sum of increases in degree in hyperconnected nodes for a group must be offset by equal, but opposite, decreases in degree for other nodes in that group. This phenomenon holds because of the mathematical relationship between average degree and graph density (Eq. 9). For simplicity, our first simulation represents the scenario where there are meaningful FC *increases* in patients for selected nodes and *unreliable* FC decreases in other nodes. This unreliability is intended to represent sampling variability that could lead to erroneous false positives.

#### Structure of whack-a-node simulation

As described above, we simulated 50 patients and 50 controls based on a groundtruth FC matrix. We estimated 100 replication samples with equal levels of noise in both groups (see Table S1). In each replication sample, we increased the mean FC in three nodes, selected at random, by *r* = 0.14. We also applied smaller decreases of *r* = −.04 to three other randomly targeted nodes^1^. This level of decrease was chosen such that group differences in analyses of nodal strength (i.e., computed on weighted graphs) were nonsignificant on average (*M p* = .19, *SD p* = .02). We applied PT to binarize graphs in each group, varying density between 5% and 25% in 1% increments. Likewise, for FC thresholding, we binarized graphs at *r* threshold between *r* = .2 and .5 in .02 increments. Finally, we retained weighted graphs for all simulated samples to estimate group differences in nodal strength. To ensure that effects were not attributable to particular nodes, we averaged group statistics across the 100 replication samples, where the targeted nodes varied randomly across samples.

In each replication sample, we estimated degree centrality for the Positive (hyperconnected), Negative (weakly hypoconnected), and Comparator nodes. Comparators were three randomly selected nodes in each sample that were not specifically modulated by the simulation. These served as a benchmark to ensure that simulations did not induce group differences in centrality for nodes not specifically targeted. To quantify the effect of PT versus FC thresholding in binary graphs, we estimated group differences (patient – control) on degree centrality using two-sample *t*-tests for each Positive, Negative, and Comparator node. For weighted analyses, we estimated group differences in strength centrality.

#### Global Insensitivity Simulation Methods: Hypothesis-to-scale matching

We simulated a dataset of 50 ‘controls’ and 50 ‘patients’ in which the fronto-parietal network (FPN) and dorsal attention network (DAN) were modulated in controls compared to the groundtruth matrix, **W**. In this simulation, the patient group had increased FC in the default mode network (DMN), a common finding in neurological disorders (Hillary et al., 2015). In controls, we increased FC strength for edges *between* FPN/DAN regions and other networks, *M r* = 0.2, *SD* = 0.1. Per-module variation in between-network FC changes was assumed to be normally distributed within each subject, *SD* = 0.1. We also increased controls’ FC on edges *within* the FPN and DAN, *M r* = 0.1, between-subjects *SD* = 0.05, within-subjects *SD* = .05. In patients, between-network FC for DMN nodes was increased, *M r* = 0.2, between-subjects *SD* = 0.1, within-subjects *SD* = 0.1. Likewise, within-network FC in the DMN was increased, *M r* = 0.1, between-subjects *SD* = .05, within-subjects *SD* = .05. That is, we applied similar levels of FC modulation to the FPN/DAN in controls and the DMN in patients, although these changes largely affected different edges in the networks between groups.

We computed the small-worldness coefficient, *σ*, according to the approach of Humphries and Gurney (2008):

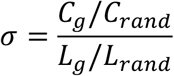

Here, *C*_*g*_ represents the transitivity of the graph, whereas *C*_*rand*_ denotes the transitivity of a random graph with an equivalent degree distribution. Likewise, *L*_*g*_ and *L*_*rand*_ represent the characteristic path length of the target graph and randomly rewired graph, respectively. Values of *σ* much larger than 1.0 correspond to a network with small-world properties. To generate statistics for equivalent random graphs, we applied a rewiring algorithm that retained the degree distribution of the graph while permuting 347,160 edges (10 permutations per edge, on average). This algorithm was applied to the target graph 100 times to generate a set of equivalent random networks. Transitivity and characteristic path length were calculated for each of these, and their averages were used in computing the small-worldness coefficient, *σ*. We also analyzed within- and between-network degree centrality for each node, *z*-scoring values within each module and density to allow for comparisons.

## Footnotes

1 Results are qualitatively similar using other values for group shifts in FC.

## Acknowledgments

We thank Zach Ceneviva, Allen Csuk, Richard Garcia, Melanie Glatz, and Riddhi Patel for their work collecting, organizing, and coding references for the literature review and manuscript.

