## Supplementary Materials for "Graph theory approaches to functional network organization in brain disorders: A critique for a brave new small-world" (from another version of this preprint: version 2, 2018-03-20)


Michael N. Hallquist

and

Frank G. Hillary

Table S1. Parameters used in network simulations

| Simulation | Parameter | Description | Value |
|---|---|---|---|
| Whack-a-node *hyper*-connectivity | $\sigma_b$ | Between-person variation in mean FC | .2 |
| | $\sigma_w$ | Within-person variation in FC | .2 |
| | $\sigma_e$ | Edge noise | .2 |
| | $\mu_{a_i}$ , $i \in P$ | Mean shift in Positive node targets in patient group | .14 |
| | $\sigma_{a_i}$ , $i \in P$ | Between-person variation in FC shifts for Positive targets | .07 |
| | $\sigma_{v_i}$ , $i \in P$ | Within-person variation in FC variation of Positive node *i* across its neighbors, *j* | .07 |
| | $\mu_{a_i}$ , $i \in N$ | Mean shift in Negative node targets in patient group | -.04 |
| | $\sigma_{a_i}$ , $i \in N$ | Between-person variation in FC shifts for Negative targets | .02 |
| | $\sigma_{v_i}$ , $i \in N$ | Within-person variation in FC variation of Positive node *i* across its neighbors, *j* | .02 |
| Whack-a-node *hypo*-connectivity | $\sigma_b$ | Between-person variation in mean FC | .2 |
| | $\sigma_w$ | Within-person variation in FC | .2 |
| | $\sigma_e$ | Edge noise | .2 |
| | $\mu_{a_i}$ , $i \in P$ | Mean shift in Positive node targets in patient group | .04 |
| | $\sigma_{a_i}$ , $i \in P$ | Between-person variation in FC shifts for Positive targets | .02 |
| | $\sigma_{v_i}$ , $i \in P$ | Within-person variation in FC variation of Positive node *i* across its neighbors, *j* | .02 |
| | $\mu_{a_i}$ , $i \in N$ | Mean shift in Negative node targets in patient group | -.14 |
| | $\sigma_{a_i}$ , $i \in N$ | Between-person variation in FC shifts for Negative targets | .07 |
| | $\sigma_{v_i}$ , $i \in N$ | Within-person variation in FC variation of Positive node *i* across its neighbors, *j* | .07 |
| Global insensitivity | $\sigma_b$ | Between-person variation in mean FC | .2 |
| | $\sigma_w$ | Within-person variation in FC | .2 |
| | $\sigma_e$ | Edge noise | .2 |
| | $\mu_{a_i}$ , $i \in \mathrm{FP, DAN}$ | Mean shift in between-module connectivity of F-P and DAN nodes in control group | .1 |
| | $\sigma_{a_i}$ , $i \in \mathrm{FP, DAN}$ | Between-person variation in between-module connectivity of FP and DAN nodes in control group | .05 |
| | $\sigma_{v_i}$ , $i \in \mathrm{FP, DAN}$ | Within-person variation in between-module connectivity of FP and DAN nodes in control group | .05 |
| | $\mu_{a_i}$ , $i \in \mathrm{FP, DAN}$ | Mean shift in within-module connectivity of F-P and DAN nodes in control group | .2 |
| | $\sigma_{a_i}$ , $i \in \mathrm{FP, DAN}$ | Between-person variation in within-module connectivity of FP and DAN nodes in control group | .1 |
| | $\sigma_{v_i}$ , $i \in \mathrm{FP, DAN}$ | Within-person variation in within-module connectivity of FP and DAN nodes in control group | .1 |
| | $\mu_{a_i}$ , $i \in \mathrm{DMN}$ | Mean shift in between-module connectivity of DMN nodes in patient group | .1 |
| | $\sigma_{a_i}$ , $i \in \mathrm{DMN}$ | Between-person variation in between-module connectivity of DMN nodes in patient group | .05 |
| | $\sigma_{v_i}$ , $i \in \mathrm{DMN}$ | Within-person variation in between-module connectivity of DMN nodes in patient group | .05 |
| | $\mu_{a_i}$ , $i \in \mathrm{DMN}$ | Mean shift in within-module connectivity of DMN nodes in patient group | .2 |
| | $\sigma_{a_i}$ , $i \in \mathrm{DMN}$ | Between-person variation in within-module connectivity of DMN nodes in patient group | .1 |

| | | |
|---|---|---|
| $\sigma_{v_i}, i \in \text{DMN}$ | Within-person variation in within-module connectivity of DMN nodes in patient group | .1 |

*Note. P* = Positive targets (3 per simulation); *N* = Negative targets (3 per simulation); FP = frontoparietal network; DAN = dorsal attention network; DMN = default mode network.

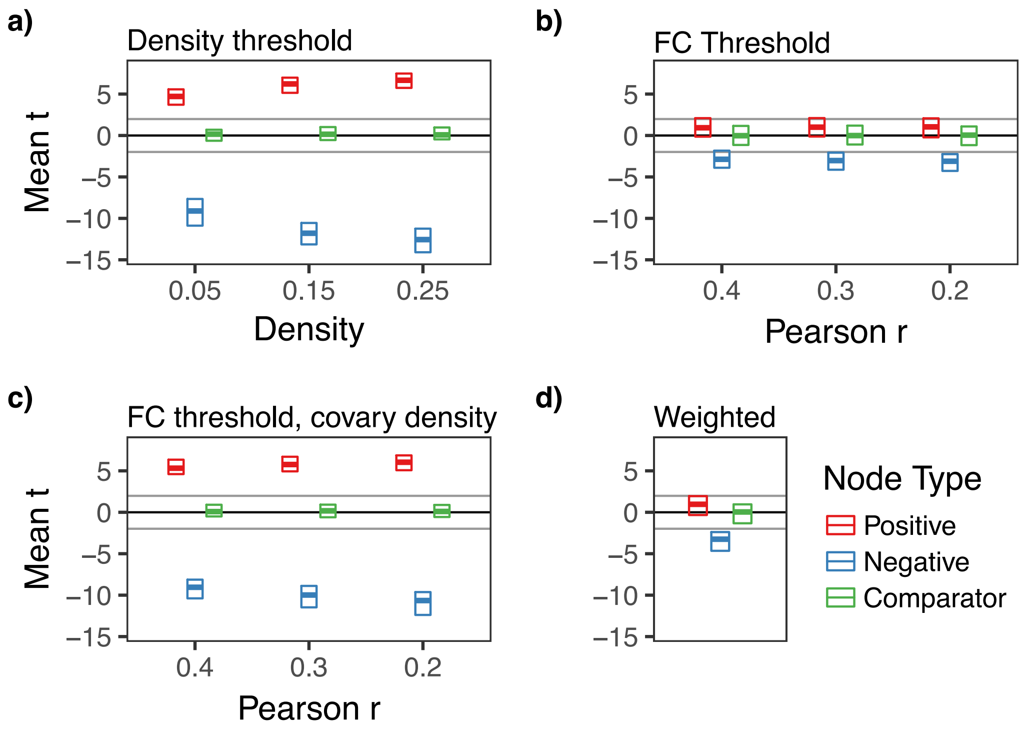


*Figure S1.* The effect of thresholding method on group differences in degree centrality when there is strong hypoconnectivity in three randomly selected nodes (Negative) and weak hyperconnectivity in three nodes (Positive). For simulation details, see Table S1, Whack-a-node hypoconnectivity. The central bar of each rectangle denotes the median $t$ statistic (patient – control) across 100 replication datasets (patient $n$ = 50, control $n$ = 50), whereas the upper and lower boundaries denote the 90$^{th}$ and 10$^{th}$ percentiles, respectively. The dark line at $t$ = 0 reflects no mean difference between groups, whereas the light gray lines at $t$ = -1.99 and 1.99 reflect group differences that would be significant at $p$ = .05. a) Graphs binarized at 5%, 15%, and 25% density. b) Graphs binarized at $r$ = {.2, .3, .4}. c) Graphs binarized at $r$ = {.2, .3, .4}, with density included as a between-subjects covariate. d) Strength centrality (weighted graphs).

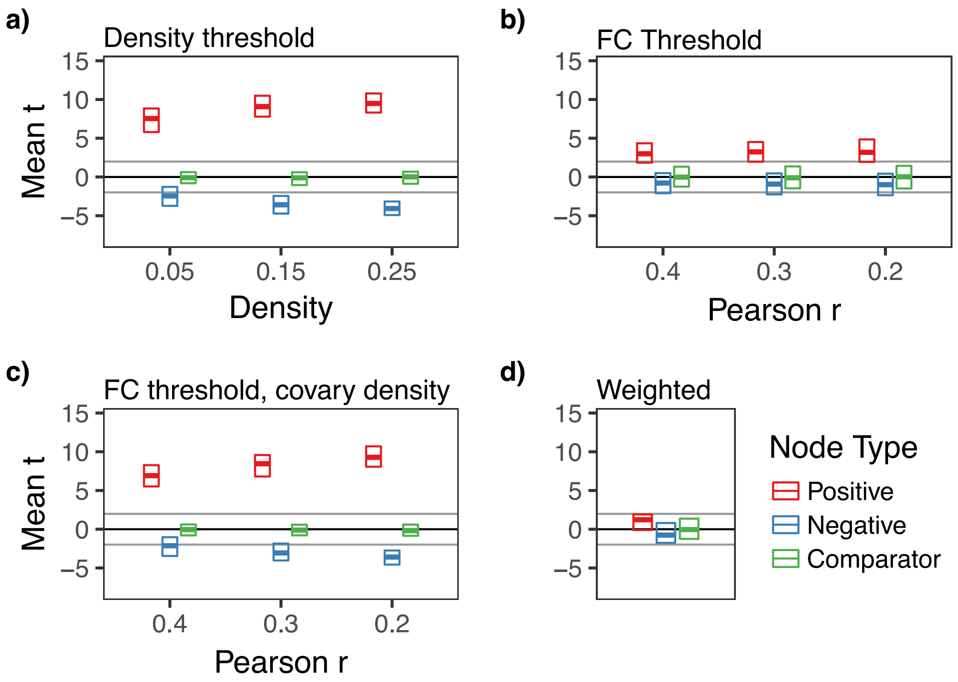


*Figure S2.* The effect of thresholding method on group differences in degree centrality when there is strong hyperconnectivity in three randomly selected nodes (Positive) and weak hypoconnectivity in three nodes (Negative). Relative to Figure 1, this simulation applied *proportional* changes to FC such that edges were shifted as a fraction of their original strength, rather than shifting edges by an absolute amount (i.e., in correlational units). Thus, FC for positive nodes was increased by 14%, on average, whereas negative nodes were decreased by 4%. All simulation parameters are identical to Table S1, Whack-a-node hyperconnectivity, but with changes applied *proportionately.*

The central bar of each rectangle denotes the median $t$ statistic (patient – control) across 100 replication datasets (patient $n$ = 50, control $n$ = 50), whereas the upper and lower boundaries denote the 90$^{th}$ and 10$^{th}$ percentiles, respectively. The dark line at $t$ = 0 reflects no mean difference between groups, whereas the light gray lines at $t$ = -1.99 and 1.99 reflect group differences that would be significant at $p$ = .05. a) Graphs binarized at 5%, 15%, and 25% density. b) Graphs binarized at $r$ = {.2, .3, .4}. c) Graphs binarized at $r$ = {.2, .3, .4}, with density included as a between-subjects covariate. d) Strength centrality (weighted graphs).
